# Chloroplast Genome Evolution, Heteroplasmy, and Inverted Repeat Dynamics in the *Elymus* Complex (Triticeae, Poaceae): Insights from Single-Molecule Sequencing of *Elymus ciliaris* and Comparative Analysis of St-Genome Lineages

**DOI:** 10.64898/2026.08.03.742459

**Authors:** Negar Karimi, Yangmeng Zhang, Hojjatollah Saeidi, Trude Schwarzacher, Qing Liu, J.S. (Pat) Heslop-Harrison

## Abstract

**Background/Objectives:** *Elymus sensu lato* (Poaceae) is arguably the largest and most complex genus in the tribe Triticeae. It includes hybrids and polyploids based on x=7 chromosomes, all including the **St** genome, forming a valuable genepool for forage grass and cereal breeding. Analysis of chloroplast genome diversity and structural dynamics is critical for resolving maternal lineages, reticulate evolution and biodiversity across this agronomically important complex, refining their taxonomy, conservation and exploitation.

**Methods:** We sequenced the complete chloroplast genome (plastome) of *Elymus ciliaris* (4x=2n=28; **StStYY** genome composition) using ultra-long Oxford Nanopore single-molecule reads and compared it to 76 additional chloroplast genomes representing major **St**-genome lineages in *Elymus s.l.* (*Pseudoroegneria* **St**; *Elymus s.s.* **StH**, **StY**; *Thinopyrum* **StJ**/**E**; *Campeiostachys* **StYH**; *Kengyilia* **StYP**). We analyzed structure, nucleotide diversity, inverted repeat (IR) dynamics, and phylogenetic signal.

**Results:** The *E. ciliaris* chloroplast genome was 135,004 bp long (38.3% GC) with a canonical quadripartite structure. Single-molecule reads (n=74) revealed heteroplasmy: two Small-Single-Copy (SSC) orientations at 30%:70% frequency, indicating an inversion polymorphism. Across the *Elymus* group, comparative analysis of chloroplast assemblies showed high structural conservation but lineage-specific IR-boundary shifts. *Kengyilia* exhibits exceptional IR expansion. Nucleotide diversity hotspots localize to the large single-copy region, especially in **StY** lineages. Phylogenies recover a monophyletic **St**-containing clade but do not delineate genera, reflecting reticulate evolution, with North American/Southeast Asian and Eurasian geographic sub-clades.

**Conclusions:** Single-molecule sequencing uncovered heteroplasmy with an inversion polymorphism in a single plant of *Elymus ciliaris*, hidden in short read assemblies. There were no other polymorphisms, as expected for chloroplast sequences (except for technical homopolymer variation). Our analyses showed that a *Pseudoroegneria*-like **St** chloroplast genome predominates as the maternal donor across *Elymus* polyploids. Variable regions and IR dynamics offer strong models for chloroplast genome evolution in reticulate lineages and suggest exploiting plastome variation to complement nuclear biodiversity studies.

## Introduction

The tribe Triticeae (Poaceae) includes the economically most important cereal crops – wheat, barley, and rye – alongside a diverse array of wild relatives that constitute a genetic reservoir for crop improvement (Dewey, 1984; Sasanuma et al., 2002; Ali et al., 2016). Among these wild relatives, the genus *Elymus* L. is arguably the largest genus within the tribe, encompassing some 186 perennial species (including hybrid species) distributed across subtropical, temperate and arctic regions worldwide (Löve, 1984; POWO, 2026). The genus is taxonomically challenging with uncertain delineation, assignment of species to genera, and definition or circumscription of species (Petersen et al., 2011; Leo et al., 2025; Lucia et al., 2019, 2020), while the evolutionary history of the species remains difficult to resolve due to polyploidy and reticulate evolution.

With a base chromosome number of x=7, *Elymus* is largely composed of allopolyploid species (Dewey, 1984; Petersen et al., 2011). Following the genomic system of classification proposed by Löve (1984), *Elymus* in the broad sense (*sensu lato, s.l.;* Melderis, 1978; 1980) comprises species with five basic genomes **St**, **Y**, **H**, **P**, and **W** (Petersen et al., 2011, Leo et al., 2025), and the proposed **J** and **E** genomes in various combinations (Pan et al., 2025). The **St** genome, derived from the diploid genus *Pseudoroegneria* (Nevski) Á. Löve (Dewey, 1967), is present in all *Elymus* species. The **H**, **P**, and **W** genomes originated from *Hordeum* L., *Agropyron* Gaertn., and *Australopyrum* (Tzvelev) Á. Löve, respectively (Dewey, 1971; Jensen, 1990; Torabinejad & Mueller, 1993), but the donor of the **Y** genome, present in the majority of Asian species, remains unidentified (Liu et al., 2006; Leo et al., 2025). In the narrow sense (*Elymus sensu stricto, s.s.*), the genus is restricted to species with **StH** and **StY** genomic constitutions, while other combinations are recognized as distinct genera: **StYH** taxa as *Campeiostachys* Drobow, **StYP** taxa as *Kengyilia* C. Yen & J. L. Yang, and species containing the **StJ/E** genome such as the genus *Thinopyrum* Á. Löve (Yen & Yang, 2013; 2022), many with reticulate evolution (Sha et al., 2025). The genomic classification requires cytogenetic confirmation, molecular validation and further integration with organelle genome data (Sha et al., 2025).

The **StY**-genome group of *Elymus* species is of particular evolutionary importance. Approximately 30 tetraploid **StY** species are restricted to temperate Asia (Liu et al., 2006), representing half of more of all known *Elymus* **StY** species (Salomon & Lu, 1992; Sha et al., 2025). Cytological analyses of artificial hybrids among **StY** species revealed that chromosome pairing gradually decreases with increasing geographical distance from parental localities, indicating substantial genetic differentiation and modification of both **St** and **Y** genomes in allopatric populations and contrasting with **StH** genomes, which maintain relatively high homology across diverse tetraploid **StH** species (Lu & von Bothmer, 1990a, b; Salomon et al., 1991; Lu & Salomon, 2004). These cytogenetic studies remain methodologically distinct from genomic approaches, and provide meticulous, labor-intensive evidence of genome differentiation that is highly informative. Despite its phylogenetic position within the enigmatic **StY** lineage, comprehensive molecular studies of the phylogenetically central species, *Elymus ciliaris* (Trin.) Tzvelev (Hu et al., 2013), a tetraploid perennial (**StStYY**; 2n=4x=28), remains limited, representing a gap in understanding **StY** genome evolution.

Chloroplast genomes (plastomes) provide an ideal system for investigating maternal lineage evolution, taxonomic revision, adaptive signatures in grasses and other species(Wu et al., 2022; Liu et al., 2020; Biswas et al., 2025; Sha et al., 2025; Chen et al., 2021; Meerprom et al., 2026), particularly in polyploid complexes. Typically, 120-160 kb in size, chloroplast genomes contain 110-130 genes encoding approximately 80 unique proteins, 30 tRNAs, and 4 rRNAs duplicated in the IRs (Palmer, 1985; Wicke et al., 2011). The typical quadripartite structure comprises two copies of inverted repeat regions (IRs, 20-28 kb) separating a large single-copy region (LSC, 80-90 kb) and a small single-copy region (SSC, 16-27 kb). Maternal inheritance in most angiosperms, including grasses, makes the chloroplast genome a powerful tool for tracing the maternal parentage of polyploid species (Salih et al., 2017).

The **St** genome’s notable ability to combine with diverse genomes and its high frequency in successful hybrids across Triticeae make the chloroplast genome particularly informative for understanding the evolutionary dynamics of this genome. By studying chloroplast variation, we can reconstruct the phylogeographic history of the **St** genome as the most frequent genome in the tribe Triticeae and elucidate maternal lineage relationships that remain obscured in nuclear phylogenies due to reticulate evolution (Mason-Gamer et al., 2010; Mason-Gamer, 2013; Hu et al. 2013).

Here we report the complete chloroplast genome sequence of a single *Elymus ciliaris* plant generated through long molecule Oxford Nanopore sequencing, which enables direct observation of full-length, single-molecule chloroplast structures and intracellular heterogeneity. We then integrate this new genome into a comprehensive comparative analysis of 77 published chloroplast genomes representing major **St**-genome lineages within the *Elymus* complex. Our specific objectives were to: (1) characterize the complete chloroplast genome structure of *E. ciliaris* with single-molecule resolution; (2) document patterns of structural variation across **St**-genome lineages; (3) identify mutation hotspots suitable for molecular marker development; and (4) reconstruct maternal phylogenetic relationships to test hypotheses about the origin and evolution of polyploid *Elymus* species.

## Materials and Methods

### Plant Material and High-Molecular-Weight DNA Extraction

Fresh leaves of a single *Elymus ciliaris* (Trin.) Tzvelev individual (USDA National Plant Germplasm System, NPGS, accession PI 632544, sometimes written PI632544; https://npgsweb.ars-grin.gov/gringlobal/search) were dark-treated for 24 hours to reduce starch content, and cut into approximately 25 mm² pieces. Approximately 2 g of leaf tissue was flash-frozen in liquid nitrogen and stored at-80°C until extraction. High-molecular-weight (HMW) genomic DNA was extracted using the Carlson Lysis Buffer protocol (Carlson et al., 1991) with modification (100 mM Tris-HCl pH 9.5, 2% CTAB, 1.4 M NaCl, 1% PEG 8000, 20 mM EDTA). Frozen leaf tissue was ground to a fine powder in liquid nitrogen using a pre-cooled mortar and pestle. The powder was divided equally between two 50 ml Falcon tubes, each mixed with 20 ml Carlson buffer (pre-warmed to 65°C) supplemented with 100 μl β-mercaptoethanol and 40 μl RNase A, then incubated at 65°C for 1 hour with gentle inversion every 30 minutes. Following cooling to room temperature, 20 ml chloroform was added to each tube, and mixtures were centrifuged at 3,500 × g for 15 minutes at 4°C. The aqueous phase was transferred to new tubes using wide-bore tips, and DNA was precipitated with 0.7 volumes of 100% (v/v) isopropanol at-80°C for 15 minutes. DNA pellets were recovered by centrifugation (3,500 × g, 45 minutes, 4°C), washed with 70% ethanol, and resuspended in G2 buffer (Blood and Cell Culture DNA Maxi Kit, QIAGEN, Hilden, Germany). Further purification employed QIAGEN Genomic-tip 500/G columns. Dissolved DNA from both tubes was pooled and passed through an equilibrated column by gravity flow. DNA was eluted with buffer QF (pre-warmed to 55°C), precipitated with 0.7 volumes of isopropanol, recovered by centrifugation, washed with 70% ethanol, air-dried briefly, and resuspended in 60 μl of TE buffer (10 mM Tris-HCl, 1 mM EDTA, pH 8.0). DNA yield and purity were assessed using Nanodrop spectrophotometry (A260/A280 = 1.86, A260/A230 = 2.12; concentration = 679 ng/μl, yielding a total of 40.74 μg HMW DNA) and agarose gel electrophoresis (Fig. 1).

**Figure 1.**
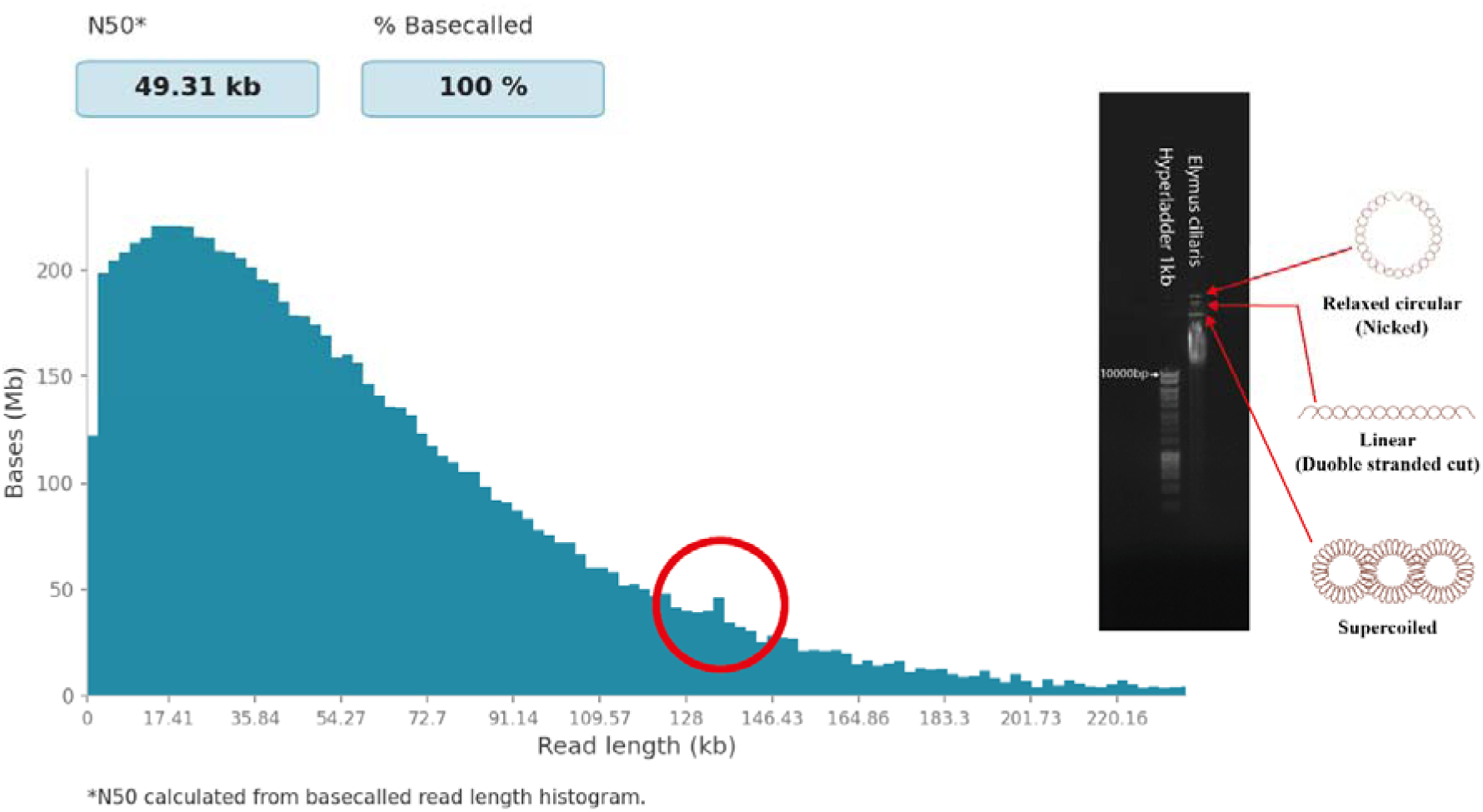
Oxford Nanopore Technology ONT single molecule read length distribution showing (red circle) the peak containing 74 near-full-length, single molecule, chloroplast reads (of the total of 280 reads between 134 and 136 kb); Inset right: Gel electrophoresis of the DNA used to construct the ultra-long sequencing library. Three large bands are visible representing nicked and relaxed circular DNA (top), linear chloroplasts, and supercoiled Covalently Closed Circle (CCC) DNA, above a smear of high molecular weight DNA >>10,000 bp.

### Nanopore Library Preparation and Sequencing

‘Ultra-long’ sequencing libraries were prepared using the Ultra-Long DNA Sequencing Kit V14 (SQK-ULK114, Oxford Nanopore Technologies, Oxford, UK) following the manufacturer’s instructions with minor modifications, most notably an increase in the amount of DNA used. HMW DNA (40.74 μg in 60 μl) was transferred to 690 μl Extraction EB (EEB). Tagmentation was performed by adding 250 μl of a mixture containing diluted Fragmentation Mix (FRA, 6 μl) and FRA dilution buffer (FDB, 244 μl) to 750 μl HMW DNA. The reaction was mixed by slow pipetting (20 times) with wide-bore tips, incubated at room temperature for 10 minutes, then at 75°C for 1 minute, followed by 5 minutes on ice. Rapid Adapter (RA, 5 μl) was added, and the reaction was incubated at room temperature for 30 minutes. Clean-up employed Precipitation Star (PS): 500 μl Precipitation Buffer (PTB) was added, mixed by rotation on a Hula Mixer (20 minutes at 3 rpm), and after confirming DNA precipitation, the supernatant was removed. The library was eluted in 300 μl Elution Buffer (EB) and incubated overnight at room temperature.

For sequencing, 90 μl of the library was mixed with 100 μl Sequencing Buffer UL (SBU) and 10 μl Loading Solution UL (LSU), incubated for 30 minutes, and loaded onto a PromethION flow cell (FLO-PRO114M, PBE41021, Oxford Nanopore Technologies) primed with a mixture of 30 μl Flush Tether UL (FTU) and 1170 μl Flow Cell Flush (FCF). Library loading employed wide-bore tips to minimize shear, with negative pressure applied via Port 2 when necessary. After 2 hours, an additional 200 μl library sequencing mix was added. The flow cell was run on a PromethION 2 Sequencing Unit Solo (PRQ-SEQ002) with MinKNOW software (version 25.05.12, https://nanoporetech.com). To increase sequence yield, the library was reloaded once after nuclease wash steps using the Flow Cell Wash Kit (EXP-WSH004; Oxford Nanopore Technologies). Raw sequencing signals were converted to fastq format using Dorado (version 7.9.8, https://nanoporetech.com) with the high-accuracy model (HAC, 400 bps). With a 99-hour runtime, 8.7 Gb of sequence data was obtained with an N50 of 47.16 kb, and reads up to 780 kb long (ONT Nanopore run report; Supplementary Data, Data S1;).

### Chloroplast Genome ‘Assembly’ and Annotation

The complete chloroplast genome of *Triticum aestivum* (NC_002762; Ogihara et al., 2002) was downloaded from NCBI as a reference. From 280 Nanopore single-molecule long reads between 134-136 kb long, 74 showed homology across their full length to the reference and were extracted for analysis. To ensure consistent starting points for alignment, all sequences were rotated to begin between the *psbA* and *rps19* genes using the Rotate tool v. 1.0 (https://github.com/richarddurbin/rotate; Durbin et al., 2023). A consensus sequence was calculated from the aligned reads (conventionally referred to as ‘assembly’, using short-reads, but here using single-molecule full-length long-reads). Gene annotation was done using the GeSeq online tool (https://chlorobox.mpimp-golm.mpg.de/geseq.html) with default parameters, employing *Triticum aestivum* (NC_002762) as reference (Tillich et al., 2017). The chloroplast genome map was generated using OGDRAW (Organellar Genome Draw version 1.3.1; https://chlorobox.mpimp-golm.mpg.de/OGDraw.html). The two consensus chloroplast genomes (with the IRa-SSC-IRb in each orientation) were deposited in NCBI GenBank under accession numbers SUB16146383 and SUB16231569 [submission number under editing via https://gb-admin@ncbi.nlm.nih.gov].

### Orientation and Repeat Structure Analysis of Single-Molecule Reads

All 74 near-full-length chloroplast reads (>134kb long) were aligned to the reference *E. ciliaris* chloroplast genome using minimap2 with default parameters at Galaxy Europe (https://usegalaxy.eu). Reads spanning the SSC region were examined for orientation relative to the reference. Dotplots of pairs of reads showing alignment over their full length in the same orientation as the reference were classified as forward, variant 1F orientation, while those with the SSC region inverted were classified as inverted, variant 1R reverse orientation. The ratio of orientations was calculated, and reads spanning the IR boundaries were examined for evidence of heterogeneous IR configurations (Palmer, 1983; Walker et al., 2015).

Repeat structure analysis employed two complementary tools. MISA (version 1.0; https://webblast.ipk-gatersleben.de/misa/) identified simple sequence repeats (SSRs) with thresholds: 10 repeats for mononucleotides, 5 for dinucleotides, 4 for trinucleotides, and 3 for tetra-, penta-, and hexanucleotides. Phobos (version 3.3.12; http://www.rub.de/ecoevo/cm/cm_phobos.htm) identified tandem repeats (7-100 bp) in imperfect search mode with repeat unit size 1-100.

### *Comparative Analysis of* Elymus sensu lato *Chloroplast Genomes*

All available complete chloroplast genome sequences of *Elymus s.l.* and related genera were downloaded from NCBI GenBank (accessed January 2026). Accessions with ambiguous annotations or possible assembly errors (missing IR, unexpected length) were excluded. The final dataset comprised 77 sequences representing: *Pseudoroegneria* (**St**), *Elymus s.s.* **StH**, *Elymus s.s.* **StY**, *Thinopyrum* (**StJ/E**), *Campeiostachys* (**StYH**), and *Kengyilia* (**StYP**). In addition 17 accessions of related genera in Poaceae and 1 accession of *Taraxacum* F. H. Wigg. as sister groups and outgroups were included in the analysis. Details of accessions used are provided in Supplementary Data Table S1.

To ensure consistency and avoid annotation bias, sequences were re-annotated using GeSeq (Tillich et al., 2017) with *Triticum aestivum* (NC_002762) as reference and default parameter settings. Multiple sequence alignments were performed using MAFFT (version 7; Katoh & Standley, 2013) with default parameters. Whole-genome synteny and structural rearrangements were visualized using Mauve (Darling et al., 2010) with a progressive alignment algorithm. Percent identity plots were generated using mVISTA (Frazer et al., 2004) in Shuffle-LAGAN mode to compare genome structures across lineages.

Nucleotide diversity (π) was calculated using DnaSP (version 6; Rozas et al., 2017) with sliding window analysis (window size = 600 bp, step size = 100 bp). Variable loci with π ≥ 0.006, a threshold where peaks were above the average level, persisted with different window sizes and corresponded to hypervariable regions in some species, were identified as potential mutation hotspots for marker development. Single nucleotide polymorphisms (SNPs) and insertions/deletions (indels) were identified from MAFFT alignments using DnaSP (version 6). Distributions across genomic regions (LSC, IRs, SSC) were calculated for each lineage. Codon usage metrics, including effective number of codons (ENC), codon bias index (CBI), and scaled chi-square (SChi²), were calculated using DnaSP (version 6) for each lineage. GC content was also determined.

Phylogenetic reconstruction included 95 sequences: 77 *Elymus s.l.* accessions, 14 related Triticeae species, and 4 outgroup sequences (*Avena* L., *Brachypodium* P. Beauv., *Taraxacum*). Maximum Likelihood (ML) analysis was done using MEGA7 (Kumar et al., 2016) with the General Time Reversible model (GTR + G + I) selected as the best-fitting model by ModelTest. Initial trees were obtained by applying Neighbor-Joining and BioNJ algorithms to pairwise distance matrices estimated using Maximum Composite Likelihood. A discrete Gamma distribution (4 categories) was used to model rate variation among sites (+G, parameter = 0.9386), with 11.88% of sites evolutionarily invariable (+I).

## Results

### Single-Molecule Resolution of Elymus ciliaris Chloroplast Genome

Ultra-long Oxford Nanopore sequencing of high molecular weight DNA from a single plant yielded 74 reads between 134-136 kb with homology to the *Triticum aestivum* chloroplast reference: near-full-length single-molecule chloroplast genomes. The complete *E. ciliaris* chloroplast genome Variant 1F was 135,004 bp long with a quadripartite structure, typical of known chloroplast genomes, comprising a large single-copy region (LSC: 1-80,626 bp), inverted repeat b (IRb: 80,627-101,437 bp, 20,810 bp), small single-copy region (SSC: 101,438-114,181 bp, 12,743 bp), and inverted repeat a (IRa: 114,182-135,004 bp, 20,811 bp) (Figure 2). Overall GC content was 38.3%. Variant 1R had the SSC in reverse, inverted, orientation between the homologous IRa and IRb. Variant 1R was 135,008 bp long, with the difference represented by extra bases in A/T homopolymers, likely to be sequencing errors.

**Figure 2.**
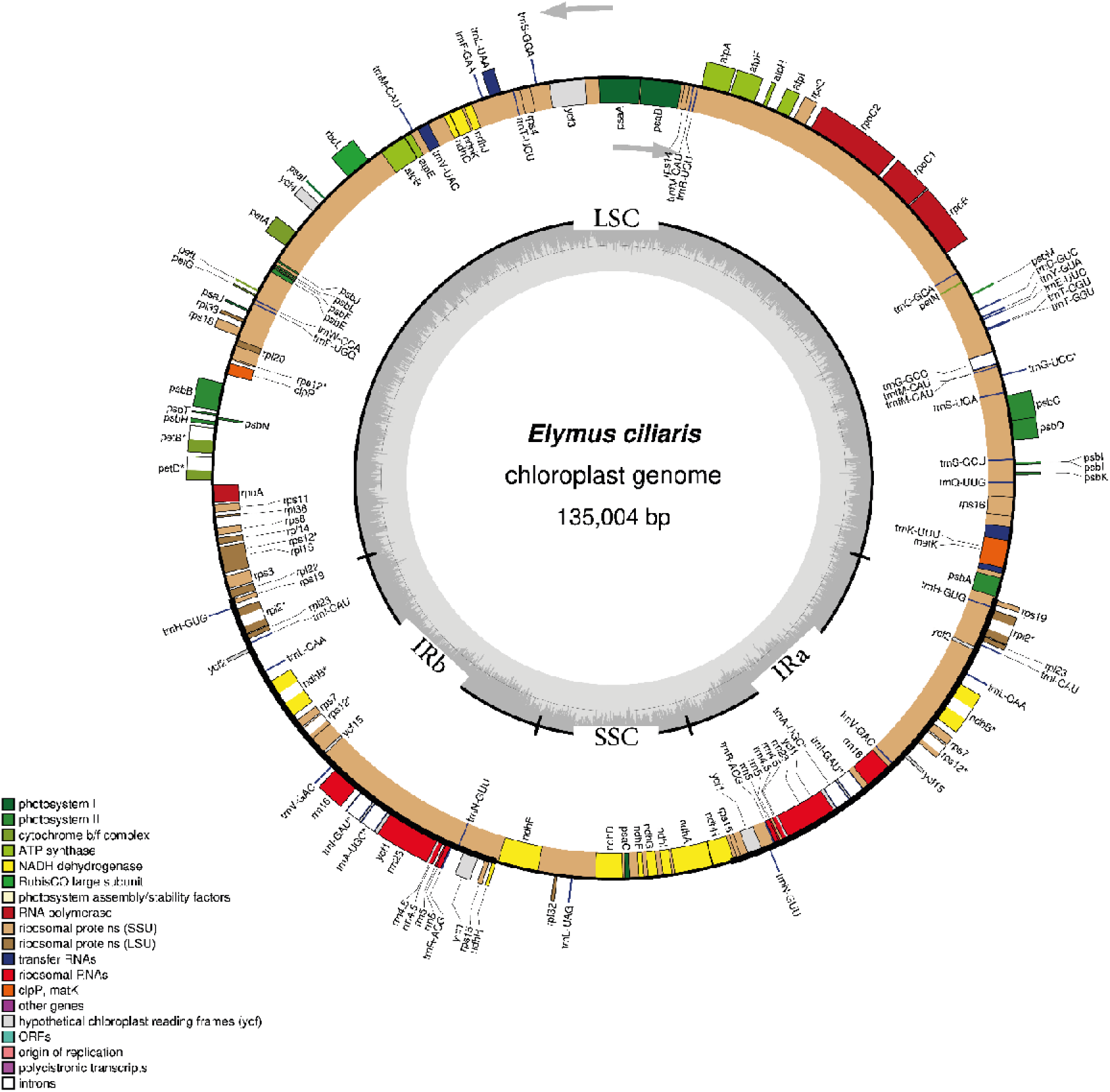
Gene map of the *Elymus ciliaris* chloroplast genome Variant 1F. Genes (total of 127) inside and outside the outer circle are transcribed clockwise and counterclockwise, respectively. Colors indicate functional groups: photosynthesis-related genes (green), transcription/translation-related genes (orange), rRNA genes (red), tRNA genes (blue). Dark gray shading inner circle represents GC content (average 38.3% GC); light gray represents AT content. IR, inverted repeat; LSC, large single copy; SSC, small single copy. See Supplementary Data Fig. S1 for Variant 1R.

### Gene content and organization

The genome contains 127 unique genes, including 82 protein-coding genes, 37 tRNA genes, and four pairs of rRNA genes (rRNA genes duplicated in IR regions). Functional categorization reveals 18 photosynthesis-related genes (photosystem I, photosystem II, ATP synthase, cytochrome b/f complex, NADH dehydrogenase) and 16 transcription/translation-related genes (RNA polymerase subunits, ribosomal proteins, translation initiation factor) (Table 1).

**Table 1.** Gene content of the *Elymus ciliaris* chloroplast genome. Numbers in parentheses indicate copy number in IR regions.

| Category | Gene Group | Gene Names |
| --- | --- | --- |
| Photosynthesis | Photosystem I | psaA, psaI, psaJ |
| Photosynthesis | Photosystem II | psbA, psbB, psbC, psbD, psbE, psbF, psbH, psbI, psbJ, psbK, psbL, psbM, psbN, psbT, psbZ |
| Photosynthesis | ATP synthase | atpA, atpB, atpE, atpF, atpH, atpI |
| Photosynthesis | Cytochrome b6/f | petA, petB, petD, petG, petL, petN |
| Photosynthesis | NADH dehydrogenase | ndhA, ndhB (2×), ndhC, ndhD, ndhE, ndhF, ndhG, ndhH, ndhI, ndhJ, ndhK |
| Photosynthesis | Rubisco | rbcL |
| Transcription /Translation | RNA polymerase | rpoA, rpoB, rpoC1, rpoC2 |
| Transcription /Translation | Ribosomal proteins (SSU) | rps2, rps3, rps4, rps7 (2×), rps8, rps11, rps12 (2×), rps14, rps15, rps16, rps18, rps19 |
| Transcription /Translation | Ribosomal proteins (LSU) | rpl2 (2×), rpl14, rpl16, rpl20, rpl22, rpl23 (2×), rpl32, rpl33, rpl36 |
| Transcription /Translation | Translation initiation | infA |
| RNA genes | Ribosomal RNA | rrn4.5 (2×), rrn5 (2×), rrn16 (2×), rrn23 (2×) |
| RNA genes | Transfer RNA | 37 tRNA genes (17 in IR, 20 in SC regions) |
| Other | Conserved ORFs | ycf1, ycf2 (2×), ycf3, ycf4 |

### LSC region organization

The LSC region contains 115 genes, beginning with *psbA* at position 92 (1,062 bp) and ending with *rpl22* at the IRb boundary. Key features include the *trnK-UUU* gene containing the *matK* intron and the *rps16* gene with its characteristic intron.

### IR region organization

Both IR regions contain identical gene sets including four rRNA genes (*rrn16*, *rrn23*, *rrn4.5*, *rrn5*), seven tRNA genes, and protein-coding genes *rpl2*, *rpl23*, *rps7*, *rps12* (3’ end), *ndhB*, and *ycf2* (Zhu et al., 2016). IRb begins with *rps19* at 80,672 bp, while IRa ends with *rps19* at 135,004 bp.

### SSC region organization

The SSC region contains 12 genes, including *ndhF*, *ndhH*, *ndhA*, *rps15*, and *ycf1*. The *ndhA* gene contains an intron, and *ycf1* spans the SSC/IRa boundary.

### Codon usage, amino acid frequency, and repeat content

Analysis of 25,514 codons revealed leucine as the most frequent amino acid (4,442 occurrences, 17.4%), encoded by six codons (UUA, UUG, CUU, CUC, CUA, CUG). Isoleucine is second most frequent (3,416 occurrences, 13.4%), encoded by AUU, AUC, and AUA. The most frequent individual codon is AAA (lysine, 1,846 occurrences), followed by UUU (phenylalanine, 1,729), AUU (isoleucine, 1,453), and AAU (asparagine, 1,402). The least frequent codon is CGC (arginine, 229 occurrences) (Supplementary Data Table S2).

MISA identified 25 simple sequence repeats (SSRs), predominantly mononucleotide A/T repeats (18 SSRs), with 9 decanucleotide repeats being most abundant. Phobos analysis revealed 703 tandem repeat units across the genome, distributed as: LSC (470 units, 67%), IRa (100 units, 14%), SSC (81 units, 12%), and IRb (52 units, 7%) (Table 2; Supplementary Table S3). Repeat unit length ranged from 7-76 bp, with hexanucleotide repeats most abundant (136 units).

**Table 2.** Number of tandem repeat units in the whole *Elymus ciliaris* chloroplast genome, large single-copy region (LSC), inverted repeat region B (IRb), small single-copy region (SSC) and inverted repeat region A (IRa).

| Repeat Unit | Whole Genome | LSC | IRb | SSC | IRa |
| --- | --- | --- | --- | --- | --- |
| Mononucleotide | 245 | 74 | 12 | 30 | 29 |
| Dinucleotide | 29 | 20 | 2 | 3 | 4 |
| Trinucleotide | 42 | 32 | 4 | 0 | 6 |
| Tetranucleotide | 68 | 29 | 8 | 15 | 16 |
| Pentanucleotide | 96 | 68 | 5 | 12 | 11 |
| Hexanucleotide | 136 | 90 | 17 | 7 | 22 |
| Heptanucleotide | 45 | 32 | 3 | 4 | 6 |
| Octanucleotide | 20 | 14 | 1 | 2 | 3 |
| Nonanucleotide | 7 | 2 | 0 | 4 | 1 |
| Decanucleotide | 1 | 0 | 0 | 1 | 0 |
| 11-mer | 3 | 1 | 0 | 1 | 1 |
| 12-mer | 0 | 0 | 0 | 0 | 0 |
| 13-mer | 1 | 0 | 0 | 1 | 0 |
| 14-mer | 3 | 1 | 0 | 1 | 1 |
| 15-mer | 2 | 2 | 0 | 0 | 0 |
| 16-mer | 1 | 1 | 0 | 0 | 0 |
| 17-mer | 0 | 0 | 0 | 0 | 0 |
| 18-mer | 0 | 0 | 0 | 0 | 0 |
| 19-mer | 1 | 1 | 0 | 0 | 0 |
| 20-mer | 2 | 2 | 0 | 0 | 0 |
| 21-mer | 1 | 1 | 0 | 0 | 0 |
| **Total** | **703** | **470** | **52** | **81** | **100** |

### Intracellular Chloroplast Heterogeneity: SSC Orientation Polymorphism

Alignment of the 74 full-length single-molecule chloroplast reads to the reference genome revealed heterogeneity in SSC orientation. Among the 74 reads, 22 (29.7%) showed the SSC region in the same orientation as the reference, while 52 (70.3%) showed the SSC region inverted relative to the reference (Fig. 3) indicating that the single *E. ciliaris* plant maintained a heterogeneous population of chloroplast genomes with two alternative configurations. Examination of reads spanning the IRa/SSC/IRb boundaries revealed that the inversion is consistent with recombination between the IR regions. All 74 reads maintained complete gene content and structural integrity, confirming that both configurations represent functional chloroplast genomes. No evidence of heterogeneity in IR configuration or LSC orientation was observed.

**Figure 3.**
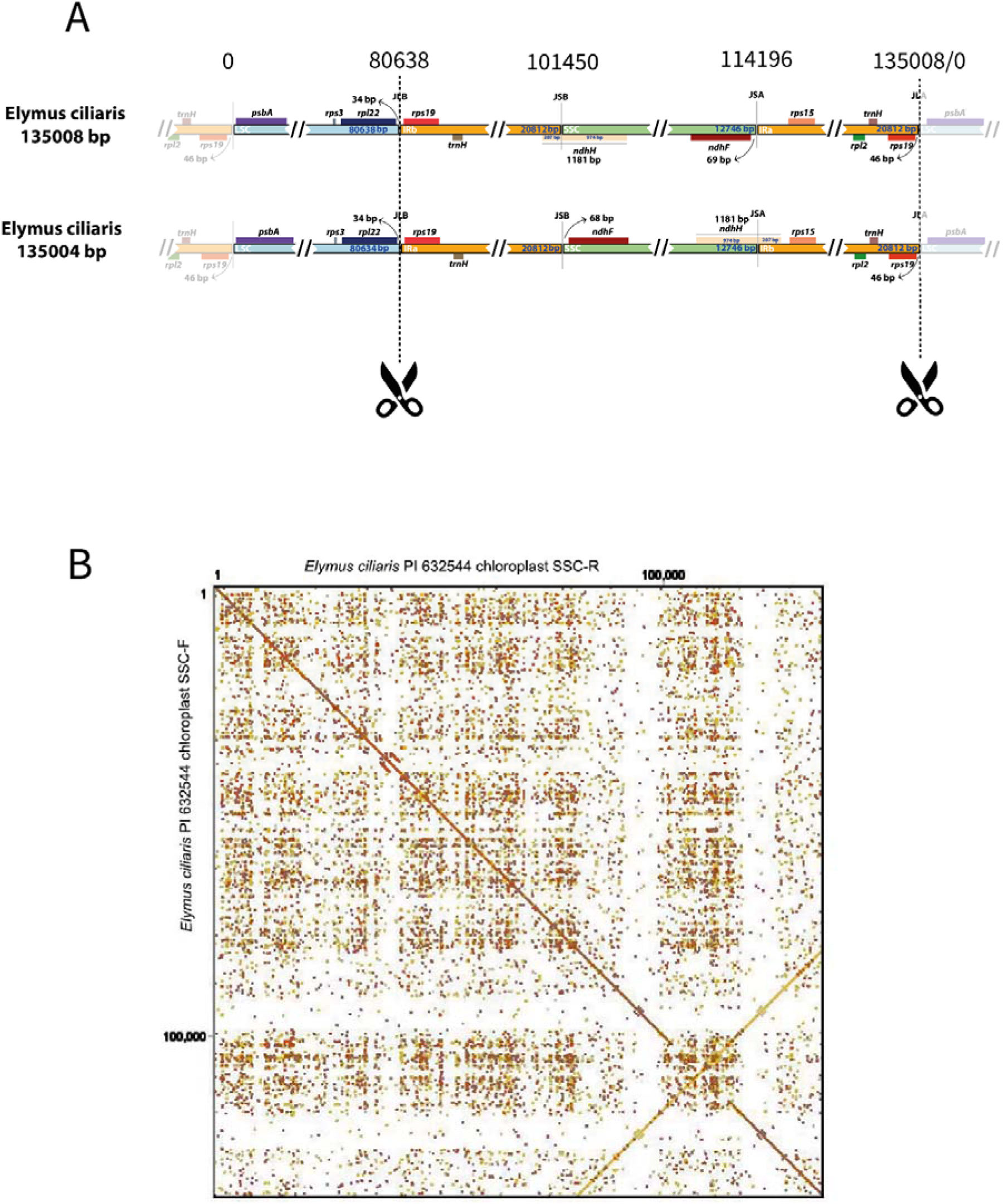
Two chloroplast isomers with inverted orientations of the small single-copy SSC region between the inverted repeats IR in *Elymus ciliaris* chloroplasts from a single plant. (A) Variant 1F orientation (22 reads, 29.7%). (B) Variant 2R, inverted, orientation (52 reads, 70.3%). The SSC region (orange) is flanked by IRa (red) and IRb (yellow). Arrows indicate directionality of SSC genes with cross indicating inversion (with ORFs on different strands). (C) Dotplot of the two isomers showing the inversion of the SSC.

### Comparative Analysis of Elymus sensu lato Chloroplast Genomes

Several features were observed while comparing 77 chloroplast genomes from six genomic *Elymus s.l.* combinations (*Pseudoroegneria* St genome, *Elymus* **StH** genomes, *Elymus* **StY**, *Thinopyrum* **StJ/E**, *Campeiostachys* **StYH**, *Kengyilia* **StYP**; Table 3, Fig. 4 and supplementary Fig. S2).

**Figure 4:**
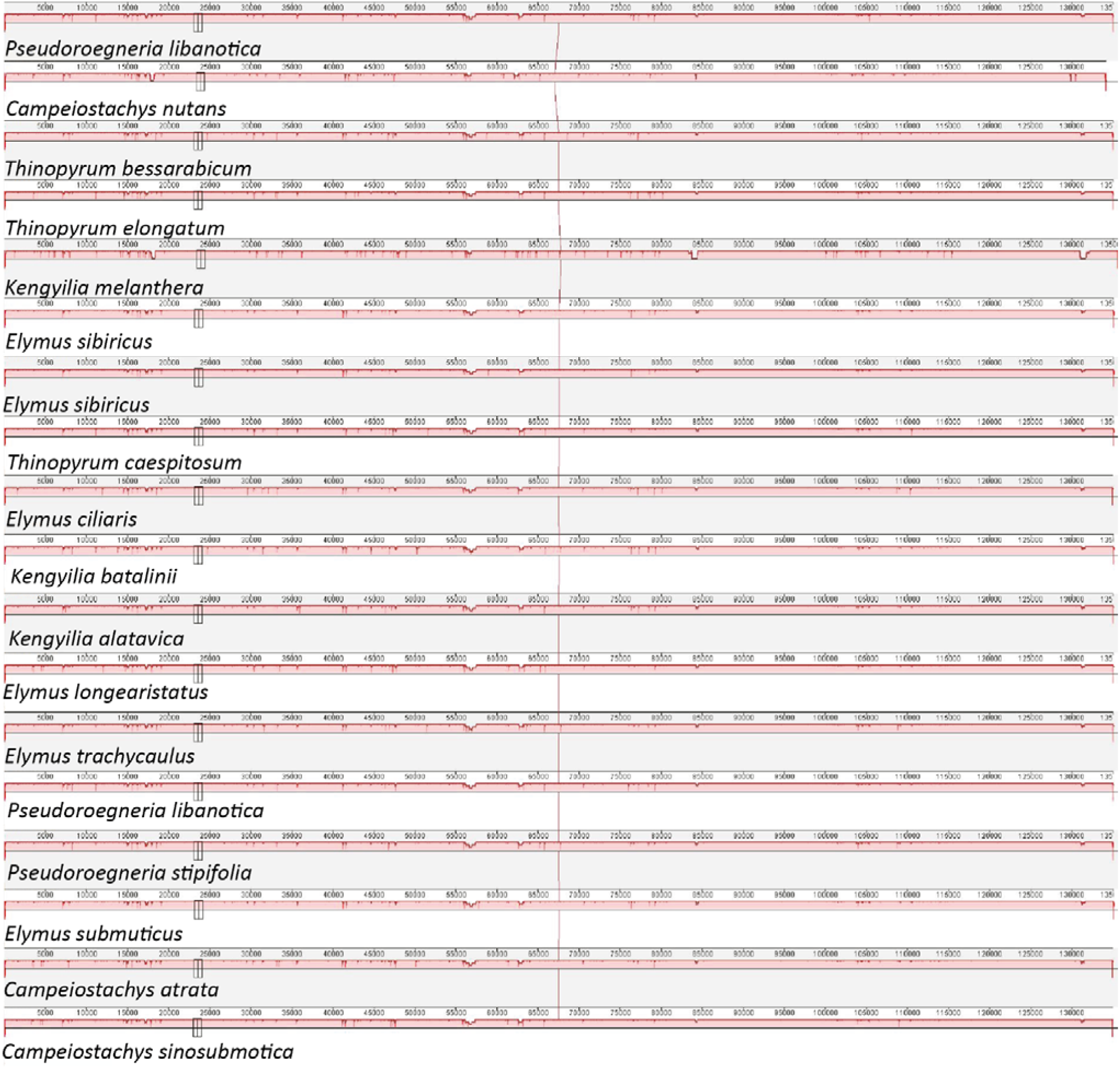
Mauve alignments comparing chloroplasts of the *Elymus s.l.* species showing nucleotide polymorphisms (red), but no notable interspecific rearrangements while reflecting small differences in length (vertical lines). Notably, no large-scale rearrangements were detected. Position in bp shown. All genomes analysed together are presented in Supplementary Data Fig. S2.

**Table 3.** Summary of Chloroplast genome characteristics across *Elymus* s.l. Lineages. Values show range (mean in parentheses) for the whole genome, the Large Single Copy region (LSC), the Inverted Repeat (IR, as one value for both IRa and IRb) and the Short Single Copy region (SSC). For details about species belonging to each lineage see Supplementary Data Table S1.

| Lineage (Genome Composition) | n <sup>1)</sup> | Whole genome Length Range (Mean) (bp) | LSC Length Range (bp) | IR Length Range (bp) | SSC Length Range (bp) | GC (%) |
| --- | --- | --- | --- | --- | --- | --- |
| *Pseudoroegneria* (**St**) | 4 | 134,972 -135,094 (135,009) | 80,566 - 80,634 | 20,813 | 12,766 - 12,767 | 38.3 |
| *Elymus* (**StH**) | 21 | 134,749 -135,460 (135,059) | 79,619 - 80,766 | 20,777 - 21,540 | 12,747 - 12,780 | 38.3 |
| *Elymus* (**StY**) | 4 | 134,972 -139,119 (135,541) | 80,557 - 80,699 | 20,813 - 20,824 | 12,752 - 12,767 | 38.3 |
| *Thinopyrum* (**StJ/E**) | 12 | 135,003 -135,094 (135,047) | 80,624 - 80,695 | 20,809 - 20,821 | 12,753 - 12,791 | 38.3 |
| *Campeiostrachys* (**StYH**) | 19 | 134,951 -135,178 (135,080) | 80,595 - 80,761 | 20,795 - 20,818 | 12,763 - 12,791 | 38.3 |
| *Kengyilia* (**StYP**) | 17 | 134,364 -135,642 (134,997) | 79,689 - 80,805 | 20,601 - 21,562 | 12,746 - 12,780 | 38.3 |
| All *Elymus* s.l. | 77 | 134,147 -139,119 (135,060) | 79,619 - 80,805 | 20,601 - 21,562 | 12,746 - 12,791 | 38.3 |

1) n = number of individuals analysed

### Genome size variation across lineages

Analysis revealed overall structural conservation with lineage-specific size variation (Table 3). Total genome length ranged from 134,147 bp (*Campeiostachys nutans* MG673520) to 135,642 bp (*Kengyilia melanthera* MH569082), with a mean of 135,060 bp. The size of the newly sequenced *E. ciliaris* (135,004 bp) falls near the mean.

### LSC region

LSC length varied from 79,619 bp (*Elymus lanceolatus* (Scribn. & J. G. Sm.) Gould) to 80,805 bp (*Kengyilia melanthera* (Keng & S. L. Chen) J. L. Yang, C. Yen & B. R. Baum) (Table 3), with most lineages showing lengths near 80,600 bp. The reduced LSC in *E. lanceolatus* and some *Kengyilia* species correlates with expanded IR regions.

### IR region

While most accessions showed the predominant IR length of 20,813 bp, notable exceptions revealed lineage-specific IR expansion/contraction events: *Elymus lanceolatus* (**StH**): IR = 21,540 bp (727 bp expansion); *Kengyilia melanthera* (**StYP**): IR = 21,562 bp (749 bp expansion); *Kengyilia hirsuta* (Keng) J. L. Yang, C. Yen & B. R. Baum (**StYP**): IR = 21,483 bp (670 bp expansion); *Kengyilia grandiglumis* (Keng) J. L. Yang, C. Yen & B. R. Baum (**StYP**): IR = 21,450 bp (637 bp expansion); *Kengyilia thoroldiana* (Oliv.) J. L. Yang, C. Yen & B. R. Baum (**StYP**): IR = 20,601 bp (212 bp contraction). These IR size variations are precisely compensated by reciprocal changes in LSC length, maintaining overall genome size within narrow limits. The expansion events are concentrated in *Kengyilia* (4/10 species show IR >21,400 bp) and one *Elymus* **StH** species, indicating lineage-specific IR boundary shifts.

### SSC region

SSC length showed minimal variation (12,746-12,791 bp), with *Thinopyrum* and*Campeiostachys* showing the highest upper limits (12,791 bp) and *Kengyilia* the lowest (12,746 bp).

### GC content

GC content was remarkably constant (38.3%) across all accessions except *Elymus repens* (L.) Gould (NC_058753, MZ169384), which showed 38.4%.

### Gene content consistency

Most genomes encoded four rRNA genes (duplicated in the IRs) and 83 tRNA genes. Exceptions included *Elymus repens* (both accessions), *E. lanceolatus*, *Kengyilia melanthera*, and *K. grandiglumis*, each containing 84 tRNA genes. Protein-coding gene (CDS) counts varied from 62-127, with the extreme low value (62 in one *E. ciliaris* accession). High-quality genomes consistently showed 118-121 CDS.

### Nucleotide Diversity and Mutation Hotspots

Nucleotide diversity (π) across all *Elymus s.l.* accessions ranged from 0 to 0.003, with a mean of 0.001. However, lineage-specific analyses revealed different diversity patterns and identified mutation hotspots with π ≥ 0.006 (Table 4; Fig. 5).

**Figure 5.**
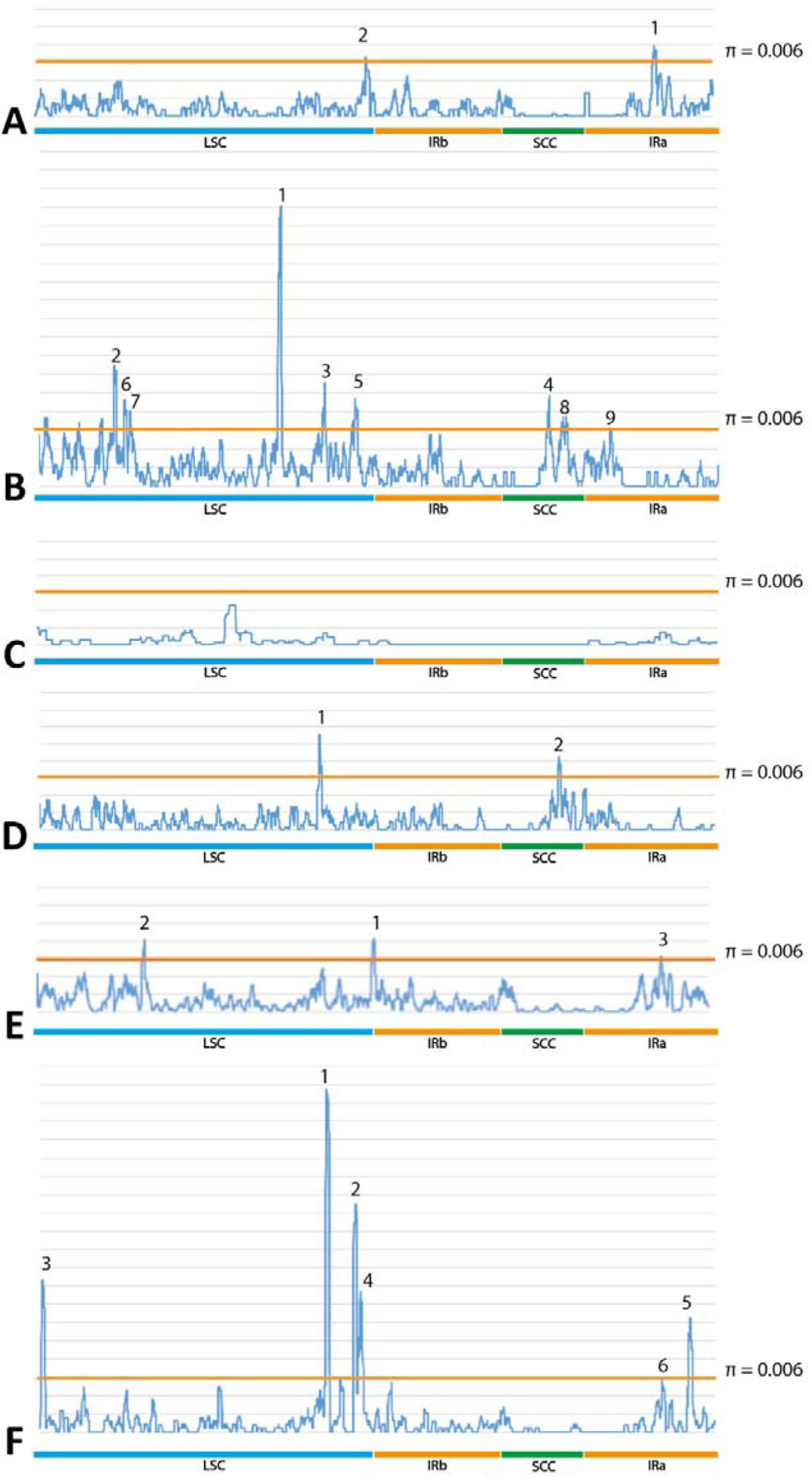
Nucleotide diversity (π) sliding window analysis across the 135kb *Elymus s.l.* lineages. Windows = 600 bp, step = 100 bp. Horizontal orange lines indicate π = 0.006 threshold for mutation hotspots. (A) *Campeiostachys* **StYH**. (B) *Kengyilia* **StYP**. (C) *Pseudoroegneria* **St**. (D) *Thinopyrum* **StJ/E**. (E) *Elymus* **StH**. (F) *Elymus* **StY**. Peaks with increased diversity are indicated as numbers 1 to 9 in each lineage. Notably, variable genomic regions are not shared between each genome combination.

**Table 4.**
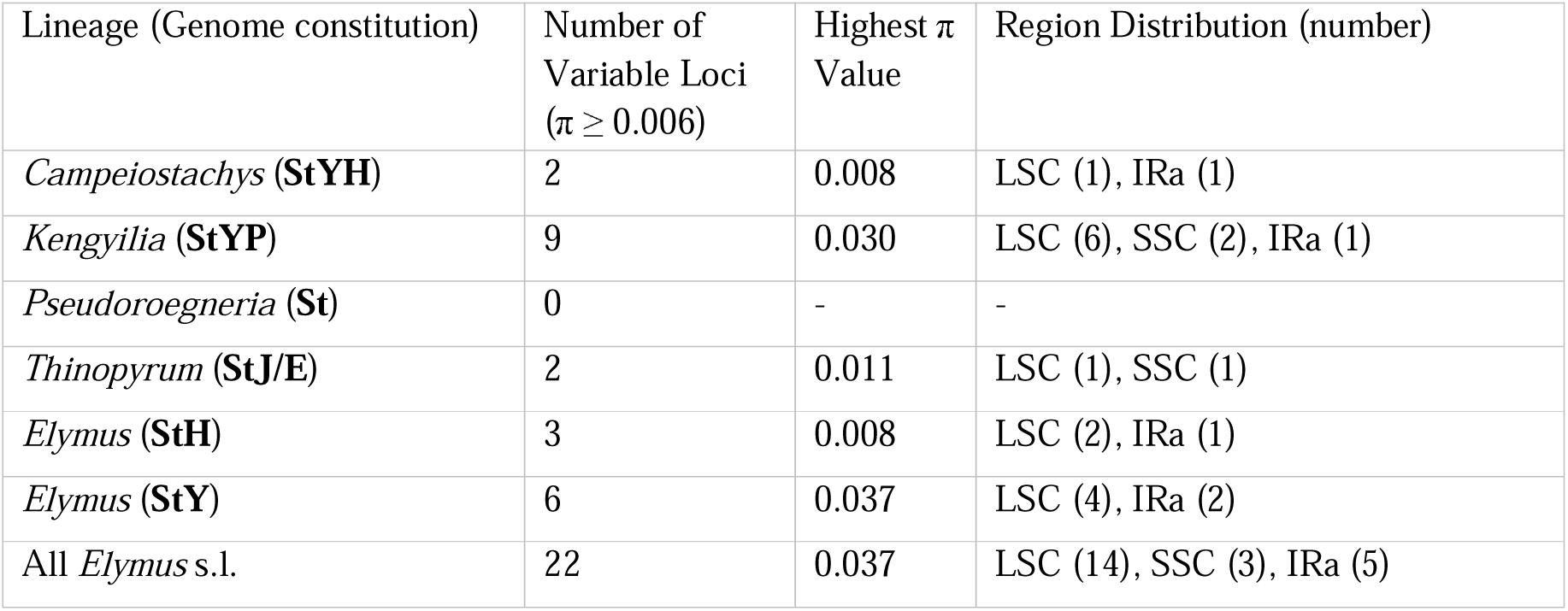
Mutation hotspots (π ≥ 0.006) across *Elymus* s.l. lineages. Large Single Copy (LSC), Short Single Copy region (SSC), Inverted Repeat (IRa)

### *Campeiostachys* StYH nucleotide diversity

Two hotspots in LSC and IRa regions were observed in *Campeiostachys* species (Fig. 5A). Peak 1 covered the length from nucleotide 125861 to 126255, mainly in the region of *rps12* gene with 0.00652<π<0.00708. Peak 2 covered the length from nucleotides 77205 to 77230, mainly in the region between the two genes of *rps8* and *rpl14*, with π=0.0066.

### *Kengyilia* StYP nucleotide diversity

The **StYP** lineage showed the most extensive variation, with nine mutation hotspots (π ≥ 0.006) distributed across different regions. Six hotspots in LSC, two hotspots in SSC and one hotspot in IRa regions were observed in *Kengyilia*species (Fig. 5B). Peak 1 covered nucleotides 48331 to 50004, with 0.00806<π<0.03012. This region contained the genes *ndhK* and *ndhC*. Peak 2 covered nucleotides 14809 to 16179, with 0.00806<π<0.03012, containing the genes *trnE*, *trnY*,and *trnD*. Peak 3 covered nucleotides 57117 to 59829, with 0.01107<π<0.00623, containing the genes *petA*, *cemA*, *pafII*, *psaL*, and *accD*. Peak 4 covered nucleotides 103640 to 104784, with 0.0061<π<0.00971, containing the genes *ndhF*, and *rpl32*. Peak 5 covers nucleotides 65254 to 66443, with 0.00721<π<0.00949, containing the genes *petL*, *petG*, and *tRNA-CCA*. Peak 6 covered nucleotides 17097 to 18510, with 0.00767<π<0.00922, containing the genes *psbM*, and *petN*. Peak 7 covered nucleotides 19904 to 20899, with 0.006<π<0.00806, containing the gene *rpoB*. Peak 8 covered nucleotides 107422 to 108431, with 0.00625<π<0.00752, containing the genes *rpl32*, and *tRNA-Leu*.

### *Pseudoroegneria* St nucleotide diversity

*Pseudoroegneria*, representing the ancestral diploid lineage, showed the lowest diversity with π value ranging from 0 to 0.005, with mean 0.0003, and with no loci exceeding the 0.006 threshold (Fig. 5C).

### *Thinopyrum* StJ/E nucleotide diversity

Two hotspots were identified in *Thinopyrum* accessions as one in LSC and one in SSC regions (Fig. 5D). Peak 1 covered nucleotides 55814 to 57018, with 0.01111<π<0.00672, containing the genes *rpl23*, and *psaI*. Peak 2 covered nucleotides 103631 to 104824, with 0.00621<π<0.00846, containing the gene *ndhF*.

### *Elymus* StH nucleotide diversity

Three variable loci were distributed across LSC, and IRa in *Elymus* **StH** accessions (Fig. 5E). Peak 1 covered nucleotides 78639 to 81072, with 0.00771<π<0.00835, containing the genes *rbcL*, *rpl23*, *accD*, and *psa1*. Peak 2 covered nucleotides 40004 to 41547, with 0.00609<π<0.00806, containing the gene *tRNA-Cys*. Peak 3 covered nucleotides 128945 to 129625, with 0.00617<π<0.00633, containing the gene *rpl32*.

### *Elymus* StY nucleotide diversity

The *Elymus* **StY** genome group showed the highest diversity peak (π = 0.037) within the LSC region, the highest value observed across all lineages (Fig. 5F). Six variable loci were identified (4 peaks in LSC region, and 2 peaks in IRa region). Peak 1 covered nucleotides 68305 to 70780, with 0.00792<π<0.03756, containing the genes *tRNA-GAA*, *ndhJ*, *ndhK*, and *ndhC*. Peak 2 covered nucleotides 73969 to 77034, with 0.00625<π<0.02494, containing the genes *atpB*, and *rbcL*. Peak 3 covered nucleotides 21238 to 22826, with 0.00667<π<0.01661, containing the genes *psbA*, and *matK*. Peak 4 covered nucleotides 76553 to 78065, with 0.00673<π<0.01548, containing the genes *rbcL*, and *rpl23*. Peak 5 covered nucleotides 130594 to 131971, with 0.00905<π<0.0125, containing the genes *ndhE*, *ndhG*, and *ndhI*. Peak 6 covered nucleotides 77480 to 78165, with 0.00598<π<0.00601, containing the genes *rpl23*, and *accD*.

### Polymorphism Distribution: SNPs and Indels

Analysis of 4297 polymorphic sites across all *Elymus s.l.* alignments revealed 1284 SNPs and numerous indels. Lineage-specific polymorphism counts varied substantially (Table 5), with *Pseudoroegneria* (1080 polymorphic sites within the lineage) and *Kengyilia* (1004 sites) showing the highest numbers despite similar sample sizes. The LSC region consistently harbored the highest proportion of polymorphisms (62-71%), followed by SSC (15-20%) and IRs (12-18%). The high indel frequency was observed in *Pseudoroegneria* (958 indels vs. 122 SNPs).

**Table 5.** Plastome polymorphism distribution across *Elymus* s.l. lineages. Large Single Copy (LSC), Short Single Copy region (SSC), Inverted Repeat (IR).

| Lineage (Genome constitution) | n <sup>1)</sup> | Total Polymorphic Sites | Number of SNPs | Number of Indels | LSC (%) | IR (%) | SSC (%) |
| --- | --- | --- | --- | --- | --- | --- | --- |
| *Pseudoroegneria* (St) | 4 | 1080 | 122 | 958 | 68 | 12 | 20 |
| *Elymus* (StH) | 21 | 392 | 128 | 264 | 62 | 18 | 20 |
| *Elymus* (StY) | 4 | 843 | 310 | 533 | 71 | 14 | 15 |
| *Thinopyrum* (StJ/E) | 12 | 626 | 242 | 384 | 64 | 16 | 20 |
| *Campeiostrachys* (StYH) | 19 | 352 | 294 | 58 | 70 | 15 | 15 |
| *Kengyilia* (StYP) | 17 | 1004 | 188 | 816 | 66 | 14 | 20 |

1) n = number of individuals analysed

### Codon Usage Patterns Across Lineages

Codon usage analysis revealed remarkable consistency across all lineages (Table 6). The effective number of codons (ENC) ranged from 55.42-56.34, values approaching 61. Codon bias index (CBI) ranged from 0.16 to 0.20, and scaled chi-square (SChi²) from 0.08 to 0.10. GC content at codon positions was identical (0.383) across all lineages. The most frequent codon across all lineages was AUU (isoleucine), contrasting with the AAA (lysine) preference observed in the single *E. ciliaris* analysis.

**Table 6.**
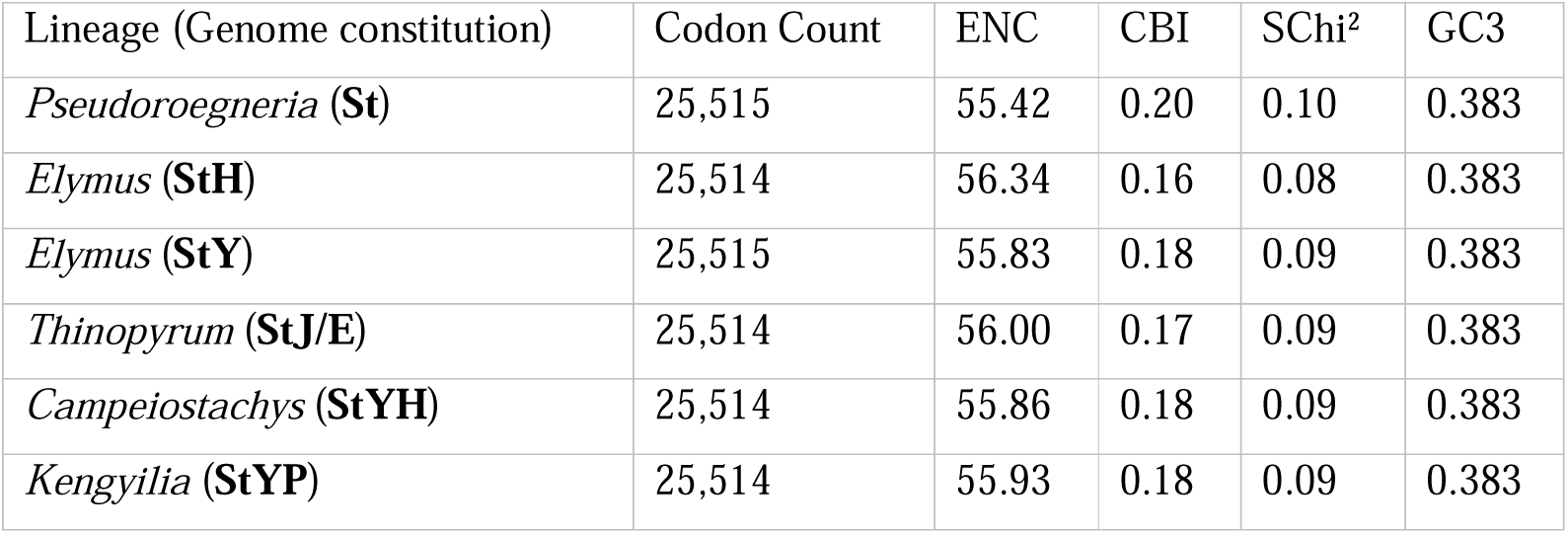
Codon usage statistics across *Elymus* s.l. Lineages. ENC, effective number of codons; CBI, codon bias index; SChi², scaled chi-square; GC3, GC content at third codon position.

| Lineage (Genome constitution) | Codon Count | ENC | CBI | SChi<sup>2</sup> | GC3 |
| --- | --- | --- | --- | --- | --- |
| *Pseudoroegneria* (St) | 25,515 | 55.42 | 0.20 | 0.10 | 0.383 |
| *Elymus* (StH) | 25,514 | 56.34 | 0.16 | 0.08 | 0.383 |
| *Elymus* (StY) | 25,515 | 55.83 | 0.18 | 0.09 | 0.383 |
| *Thinopyrum* (StJ/E) | 25,514 | 56.00 | 0.17 | 0.09 | 0.383 |
| *Campeiostrachys* (StYH) | 25,514 | 55.86 | 0.18 | 0.09 | 0.383 |
| *Kengyilia* (StYP) | 25,514 | 55.93 | 0.18 | 0.09 | 0.383 |

### Phylogenetic Relationships and Maternal Lineage Evolution

Maximum Likelihood phylogenetic analysis of 95 sequences produced a tree (log likelihood = - 97,815.33) (Fig. 6) with six main clades. among which the first five clades contained sister taxa used in the analysis. Among them clades C, D, and E are non-St genome Triticeae members. The fifth clade labeled as **St** clade (Fig. 6) contained all *Elymus s.l*. Species with St genome. This clade was further divided into two sub-clades each containing different *Elymus s.l.* species from different geographical origins. Clade I contained North American and Southeast Asian *Elymus* species, while clade II contained *Elymus* species from Central Asia toward Europe. Subsequent divisions showed correlation in part with genome compositions.Three *Kengyilia melanthera* species fell out of the **St** Clade.

**Figure 6.**
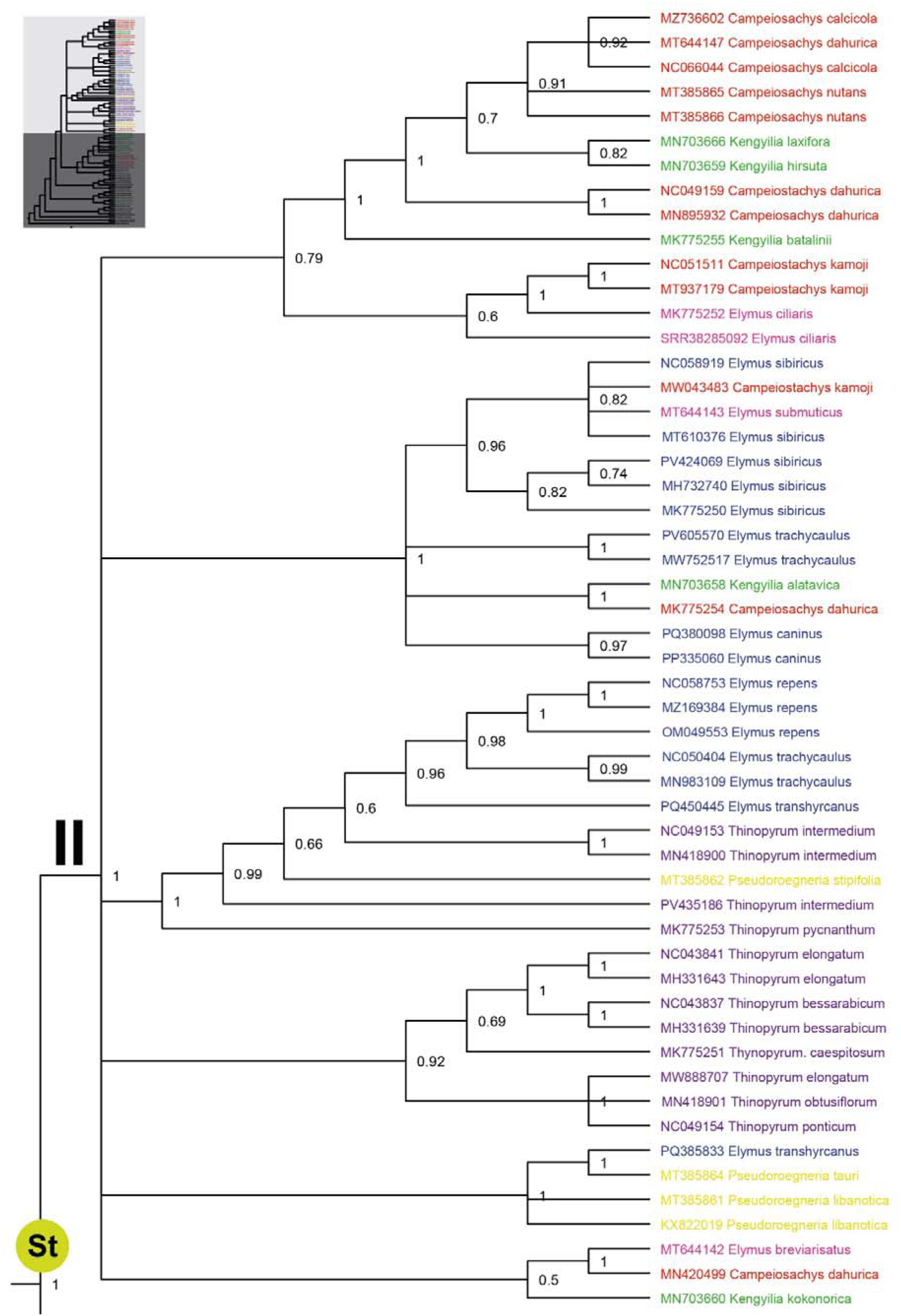
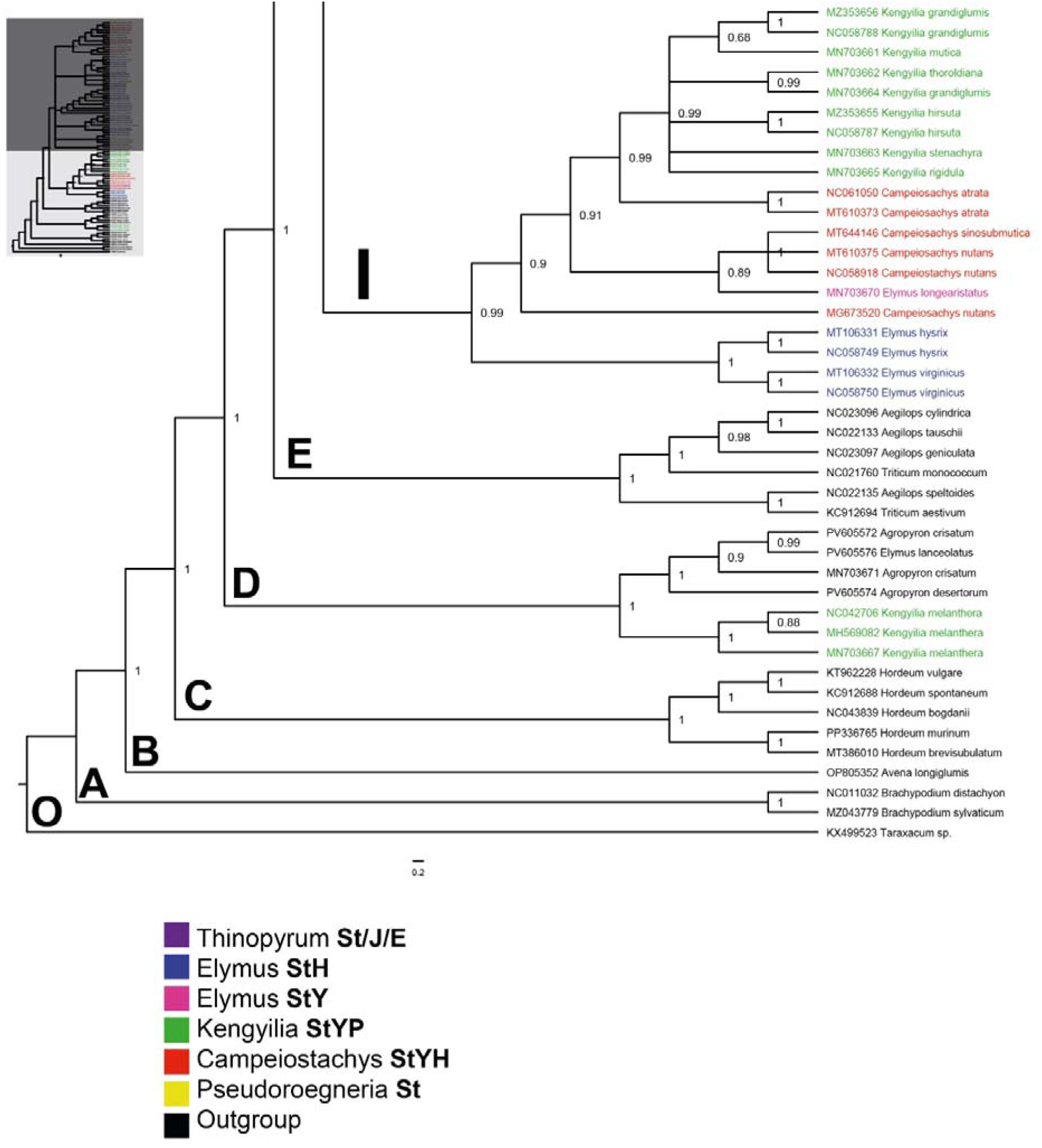
Maximum Likelihood phylogeny of *Elymus s.l.* and other representative Triticeae based on complete chloroplast genomes. The tree shows relationships among 95 accessions, with *Elymus s.l.* forming a monophyletic group labeled as St in green circle. Colors indicate genomic constitution: *Pseudoroegneria* (**St**, yellow), *Elymus* **StH** (blue), *Elymus* **StY** (pink), *Thinopyrum* (**StJ/E**, purple), *Campeiostachys* (**StYH**, red), *Kengyilia* (**StYP**, green), outgroups (black). Clade I (North American and Southeast Asian lineages) and Clade II (Eurasian lineages) are indicated. Values above branches indicate ML bootstrap support; values >70% are regarded as well-supported.

## Discussion

### Single-Molecule Resolution Reveals Intracellular Chloroplast Heterogeneity

The application of ultra-long Oxford Nanopore (Fig. 1) sequencing to *Elymus ciliaris* provides unprecedented resolution of chloroplast genome structure (Fig. 2) at the single-molecule level. Among 74 full-length chloroplast reads, from a single plant, we observed two alternative orientations of the small single-copy region in a 22:52 (1:2.4; 30%:70%) ratio (Fig. 3), demonstrating that individual plants maintain a heterogeneous population of chloroplast genomes. This phenomenon - chloroplast heteroplasmy mediated by IR recombination - has been documented in several plant species (Palmer, 1983; Walker et al., 2015; Zhang et al., 2025); our single-molecule approach provides direct quantification of the two isoforms in a natural population of chloroplasts in a single plant. However, conventional assembly algorithms with massively parallel sequences would not detect the inversion because it is flanked by inverted repeats, so the occurrence of heterogeneity in the SSC orientation is rarely reported.

The 1:2.4 ratio of reference:inverted orientation suggests that both configurations are functional and coexist stably within the cell or individual. The inversion mechanism involves homologous recombination between the IR regions, which flank the SSC as direct repeats (Palmer, 1983). This recombination flips the SSC orientation, with a characteristic region of micro-homology (11 base pairs; Fig. 3) without altering gene content or order, producing two equally viable, functional, chloroplast genome configurations. Notably, we did not find any significant heterogeneity outside the inversion, in agreement with most plastid assembly studies (outside the species with biparental inheritance; Dominicus et al., 2025) showing the absence of heteroplasmy at the SNP level (Wicke et al. 2011). Thus the likely frequent homologous recombination and gene conversion which homogenizes sequences across copies (‘copy correction’; Ruhlman et al., 2017) is not affected by the presence of the inversion.

The biological significance of the inversion heterogeneity remains unclear. It may represent neutral variation generated by recombination frequency, and the two isoforms have different replication or expression properties that maintain their coexistence. Polymorphisms elsewhere in the chloroplast increase genetic diversity and may facilitate adaptation or maintaining the functional stability of functional genes, balancing genetic stability and flexibility in chloroplast genome evolution (Wang et al., 2008). However, it seems unlikely that the presence of two SSC orientations resulting from recombination have an adaptive or functional consequence given the genes remain in the SSC and LSC; the orientations are a consequence of the presence of homologous recombination mechanisms in the chloroplast genome. Whether there is a replication advantage or higher frequency of recombination producing the predominance of one orientation (70%) is unclear. Similar observations in *Triticum* L. (Zhang et al., 2025) indicate that the phenomenon may be widespread in grasses.

This finding has practical implications for chloroplast genome assembly and analysis. Standard assembly approaches that assume a single homogeneous genome, will mask the SSC orientation heterogeneity or produce chimeric assemblies. Researchers should be aware that published chloroplast genomes may represent one isoform from a heterogeneous population, and comparative analyses should consider orientation. There is no evidence of the SSC inversion polymorphisms being useful for phylogenetic analysis.

### The Dynamic Nature of Inverted Repeat Boundaries

Our comparative analysis of chloroplast genomes from 77 species reveals that while overall structure is highly conserved (Fig. 4), the inverted repeat regions exhibit lineage-specific expansion and contraction events (Table 3). The predominant IR length of 20,813 bp in most accessions could be taken to represent the ancestral state, but several lineages show significant deviations. The most dramatic expansions occur in *Kengyilia* (**StYP**), where four species show IR lengths >21,400 bp - expansions of 637-749 bp relative to the ancestral state. *Elymus lanceolatus* (**StH**) shows a similar expansion (21,540 bp). These expansions are precisely compensated by reciprocal contractions in LSC length, maintaining overall genome size within narrow limits.

IR expansion/contraction is a well-documented mechanism of chloroplast genome evolution (Edlund et al., 2025; Goulding et al., 1996; Zhu et al., 2016). The IR boundaries are dynamic regions where gene conversion and recombination can shift the IR into adjacent single-copy regions, duplicating genes in the process (Wang et al., 2008). The concentration of expansion events in *Kengyilia*, a genus of high-altitude Central Asian grasses, suggests either lineage-specific mutation rates, or selective pressures, favouring expanded IRs. Expanded IRs can increase the copy number of genes at the boundaries, for example in the hemiparasitic order Santalales (Edlund et al., 2025), potentially affecting expression levels. In *Kengyilia melanthera* (IR = 21,562 bp), the expansion has duplicated portions of *ycf1* and *ndhF* genes involved in chloroplast function and stress response (Drescher et al., 2000). Whether this duplication confers adaptive advantages in high-altitude environments requires further investigation. The observation that *Pseudoroegneria* (the ancestral **St** diploid) maintains the ancestral IR length (20,813 bp) while its polyploid descendants show expansions suggests that polyploidization may facilitate selective constraints on IR boundaries or increase the frequency of recombination (Zhang et al., 2025).

### Mutation Hotspots: Tools for Phylogenetics and Population Genetics

The identification of 22 mutation hotspots (π ≥ 0.006) across *Elymus s.l.,* in line with peaks found in other genera (Dong et al., 2012), provides valuable tools for genotyping, population and evolutionary studies. The concentration of hotspots in LSC (61%) confirms this region as the fastest-evolving part of the chloroplast genome, consistent with studies across angiosperms (Shaw et al., 2007). The presence of SSC hotspots in *Kengyilia* and *Thinopyrum* (but not other lineages) indicates lineage-specific shifts in selective constraints or mutation rates in these taxa.

The exceptional diversity in **StY** lineages (π = 0.037 in LSC) is particularly noteworthy. The Asian **StY** genome group has long been recognized as a cytogenetically diverse group (see Introduction). Our results show that this diversity extends to the chloroplast genome, suggesting ancient divergence of maternal lineages or accelerated mutation rates in this group. The *Kengyilia* hotspots, particularly the three SSC loci, are unusual, as the SSC is typically the more conserved region (Shaw et al., 2007). Their presence suggests that *Kengyilia* chloroplasts have experienced different evolutionary dynamics than other **St** lineages, possibly related to their high-altitude distribution or unique genomic constitution (**StYP**). This high chloroplast diversity suggests there may be useful variation to use in forage grass and cereal breeding. The absence of mutation hotspots in *Pseudoroegneria* (π consistently <0.006) contrasts with its high overall polymorphism count (1080 sites). This pattern indicates that variation in *Pseudoroegneria* is dispersed across the genome rather than concentrated in hotspots, consistent with its status as an ancestral diploid lineage.

### Codon Usage: Evidence for Uniform Selective Constraints

The consistency of codon usage metrics across all lineages, despite differences in ploidy, genomic constitution, and geographic distribution, provides evidence that chloroplast genes probably experience similar selective constraints across the *Elymus* complex. ENC values (55.4–56.3) indicate weak codon bias, characteristic of genes with relatively low expression levels and limited translational selection (Morton, 2003; Suzuki & Morton, 2016; Mazumdar et al., 2017; Salih et al., 2017). The identical overall GC content (38.3%) and GC3 (38.3%) across lineages suggest that mutational pressures acting on chloroplast genes have been stable since the divergence of these lineages, and contrast with the concept that codon usage is driven by lineage specific mutational biases or strong GC biased gene conversion (gBGC), instead pointing to conserved selective constraints and/or a shared mutational landscape in *Elymus* chloroplasts. This interpretation is supported by broader chloroplast genome studies (e.g. Mazumdar et al., 2017; Liu et al., 2020; Biswas et al., 2026): chloroplast genomes commonly show low overall codon bias, strong A/T preference at third codon positions (Liu et al., 2020, in *Avena*: AT3 c. 70%), widespread purifying selection on protein coding genes, and associations of repeats/indels with local polymorphism, consistent with an AT biased mutational environment and limited recombination/gBGC effects in chloroplast genomes, while strong selection of species would lead to lineage-specific domestication signatures (Shah et al., 2026).

The shift from AAA (lysine) as the most frequent codon in *E. ciliaris* to AUU (isoleucine) in the multi species analysis shows that single genome analyses can be influenced by species specific properties that average out in comparative studies. The AUU preference across the complex may reflect the ancestral codon usage pattern in Triticeae chloroplasts.

### Phylogenetic Insights: The **St** Genome as Unifying Element and the Challenge of Reticulation

The phylogenetic analysis provides both confirmation and complication of our understanding of *Elymus* evolution. The strong support for monophyly of all **St-**containing lineages confirms that the **St** genome, derived from *Pseudoroegneria*-like ancestors, is the unifying element in this polyploid complex. This supports the classification system by Melderis (1978), which considered all **St** containing species as *Elymus s.l.* Chloroplast phylogeny cannot recover the reticulate evolution nor resolve genera defined by genomic constitution (Löve, 1984; Dewey, 1984; Yen & Yang, 2013), but shows **St**-lineage species clusters. The mixture of taxa with different genomic constitutions (**StH, StY, StJ/E, StYH, StYP**) within the same chloroplast clades indicates that polyploidization events have repeatedly involved the same or closely related maternal donors. Secondly, the distribution of *Elymus* **StH** species across both Clade I and Clade II suggests that **StH** tetraploids have originated multiple times from different *Pseudoroegneria* × *Hordeum* hybridization events (Leo et al., 2025). Similarly, the presence of *Kengyilia* in both clades, and even some out of **St** clade, indicates that **StYP** hexaploids have multiple independent origins (Chen et al., 2021). Populations with the same genomic constitution may have different evolutionary histories and potentially different traits.

### Marker Development and Future Directions

Using the nucleotide diversity analysis (Fig. 5), primer pairs targeting mutation hotspots with high π value (Supplementary Table S4) can be designed to target mutation hotspots (as in other genera: eg Biswas et al., 2026). These may be valuable for phylogeographic studies (tracing maternal lineages of *Elymus* populations across their geographic ranges) and species delimitation, feeding into conservation genetics and crop wild relative research by characterizing the diversity of chloroplast haplotypes for introgression into wheat and barley. Although relaxed selective constraints in non-coding DNA means that intergenic regions may be more polymorphic compared to coding regions, there is the potential to exploit adaptive genes (either with direct or linked markers) in new lineages. Wang et al. (2000) in an experiment over many decades, have shown the effect of chloroplast genome exchanges in wheat lines, and adaptation or resistances may be important selective characters.

Future work should integrate these chloroplast data with nuclear genome markers (e.g., from low-copy nuclear genes, RAD-seq, or whole-genome sequencing) to fully resolve the complex evolutionary history of the *Elymus* polyploid complex. The combination of maternally inherited chloroplast markers and biparentally inherited nuclear markers will enable reconstruction of both maternal and paternal contributions to polyploid genomes, revealing the full reticulate history that single-genome systems cannot capture (Mason-Gamer et al., 2010). Importantly, the results have implications for chloroplast genome evolution in reticulate plant groups and for exploiting wild relatives of cereals and increasing diversity available in plastomes.

## Conclusions

This study provides a comprehensive analysis of chloroplast genome evolution and diversity in *Elymus* complex. Key findings include: 1) Single-molecule resolution of the *E. ciliaris* chloroplast genome revealed intracellular heterogeneity in SSC orientation (30%:70% ratio), demonstrating that plants maintain mixed populations of chloroplast isoforms. 2) Structural conservation across 77 genomes confirms the stability of chloroplast architecture, while IR dynamics, particularly expansions in *Kengyilia* (up to 21,562 bp), reveal lineage-specific boundary shifts. 3) Mutation hotspots (22 loci with π ≥ 0.006) provide robust molecular markers, with **StY** lineages showing the highest diversity (π = 0.037) and *Kengyilia* showing unique SSC variation. 4) Phylogenetic analysis confirms **St**-genome monophyly but fails to resolve generic boundaries within *Elymus* s*.l.*, indicating extensive chloroplast capture and multiple origins of polyploid taxa with the same genomic constitution. These findings advance our understanding of chloroplast genome evolution in polyploid complexes, provide practical tools for future research, and highlight the necessity of multi-genome approaches for resolving reticulate evolutionary histories.

## Supplementary Data

Supplementary data are available and consist of the following:

**Supplementary Figure S1**: The maps of the chloroplast variant 1R with the SSC between IRa and IRb inverted compared to 1F (Figure 1). **Figure S2:** Mauve alignment for each group of species. **Table S1.** Complete list of GenBank accessions used in this study. **Table S2.** Codon usage table for *Elymus ciliaris*. **Table S3.** Simple sequence repeats (SSRs) identified in the chloroplast genome. **Table S4.** Primer sequences targeting mutation hotspots. **Data S1.** Nanopore sequencing run report for flow cell PBE41021. As reported in the Materials and Methods, the POD5 file created during the run was re-called with a later Dorado software version (7.9.8) with higher-accuracy parameters for the analysis presented; many homopolymers were resolved and some reads were longer.

## Author Contributions

NK: conceptualization, experimental work, data analysis, manuscript drafting; HS: conceptualization, supervision, data interpretation, manuscript revision; PHH: conceptualization, supervision, funding acquisition, manuscript revision; YZ: DNA extraction, Nanopore sequencing and analysis; TS: supervision, methodology, manuscript revision; QL: comparative analysis, funding acquisition, methodology, phylogenetic reconstruction.

## Data Availability Statement

The complete chloroplast genome sequence of *Elymus ciliaris* has been deposited in NCBI GenBank accession number [currently being edited by NCBI under Submission numbers SUB16146383 ECI_cpDNA_SSC_reference_orientation PZ609788 and SUB16146383 ECI_cpDNA_SSC_inverted_orientation PZ609789. Nanopore fastq files from *Elymus ciliaris* are uploaded to SRA Sequence Read Archive at NCBI under project PRJNA1458227 https://www.ncbi.nlm.nih.gov/sra/?term=PRJNA1458227. The terabyte size ONT POD5 files are available on request from the authors. Under the Fort Lauderdale Agreement (see Foster & Shard, 2007), these data are released early for community use with the expectation that users will acknowledge the data generators and consult them before undertaking or publishing large-scale genome-wide analyses; the authors are currently pursuing such analyses.

## Supporting information

Karimi_Elymus_SupplementaryTable S2

Karimi_Elymus_SupplementaryTable S3

Karimi_Elymus_SupplementaryTable S4

Karimi_Elymus_SupplementaryTable Data S1

Karimi_Elymus_SupplementaryFigureS1

Karimi_Elymus_SupplementaryFigureS2

Karimi_Elymus_SupplementaryTable S1

## Acknowledgments

We thank the USDA National Plant Germplasm System (NPGS) for providing seed material (PI 632544). Nanopore sequencing was performed at the University of Leicester Genomics Facility with thanks to Dr Nic Sylvius and Professor James Higgins. We are grateful to Dr Orie Shaw, Oxford Nanopore, for assistance with the PromethION system and methods.

