## Supplementary material for "Chloroplast Genome Evolution, Heteroplasmy, and Inverted Repeat Dynamics in the *Elymus* Complex (Triticeae, Poaceae): Insights from Single-Molecule Sequencing of *Elymus ciliaris* and Comparative Analysis of St-Genome Lineages": Karimi_Elymus_SupplementaryTable S4

**Supplementary Table S4**. Primer sequences targeting mutation hotspots or heterozygous regions in chloroplasts. The markers can be used to identify regions characteristic of each chloroplast genome.

| Genome variability | Name of primer | Sequence | Product Length | GC Content | Melting Temperature (Tm) |
| --- | --- | --- | --- | --- | --- |
| StYH | Ely-Rps12-F | GGATAGGGATAGAGGAAGAGGGAA | 395 | 50.0 | 60.0 |
| StYH | Ely-rps12-R | TTTTCACCTCATACGGCTCC |  | 50.0 | 57.6 |
| StYP | Ely-ndhj-F | ACAGACGGAATTCTAGGATT | 378 | 40.0 | 53.4 |
| StYP | Ely-ndhj-R | AAGAAAAAGCAGGGCAATC |  | 42.1 | 54.1 |
| StYP | Ely-ndhK-F1 | AAGGTACTGGACTTTTGG | 302 | 44.4 | 51.2 |
| StYP | Ely-ndhK-R1 | CTATAGTACTGTTCGGGGAGTTGAC |  | 48.0 | 60.2 |
| StYP | Ely-ndhK-F2 | TTCCCCCTGTAATAGTACA | 412 | 42.1 | 51.5 |
| StYP | Ely-ndhK-R2 | AGGAGCCTTGGAATGGTCTT |  | 50.0 | 58.6 |
| StYP | Ely-ndhC-F | TCGATAAAAACGGATACACC | 432 | 40.0 | 53.2 |
| StYP | Ely-ndhC-R | ACAAAGGCTAAACTAAGCGC |  | 45.0 | 56.7 |
| StYP | Ely-trnA-F1 | GGAAAAAGTCTTTGCTTTGG | 140 | 40.0 | 53.5 |
| StYP | Ely-trnA-R1 | ACTAGACGATAGGGGCGT | 140 | 45.8 | 56.7 |
| StYP | Ely-trnA-F2 | GGACATCTCTCTTTCAAGGAGG | 699 | 50.0 | 57.9 |
| StYP | Ely-trnA-R2 | TGATCTTGCTCCCTGCCC | 699 | 61.1 | 59.0 |
