## Supplementary material for "Chloroplast Genome Evolution, Heteroplasmy, and Inverted Repeat Dynamics in the *Elymus* Complex (Triticeae, Poaceae): Insights from Single-Molecule Sequencing of *Elymus ciliaris* and Comparative Analysis of St-Genome Lineages": Karimi_Elymus_SupplementaryTable Data S1

### PromethION 2 Solo P2S-02009-A Sequencing final run report

Oxford Nanopore Technologies

25 Sept 25, 11:54 UTC+1:00 — 29 Sept 25, 14:58 UTC+1:00 · ElymusCiliaris-YM2 · ECI_YM2 · P2S-02009-A
Protocol run ID: 3d5de01d-bcd0-4667-9d2c-a39cd63ccdb5

Run summary | Run configuration | Sequence output | Run health | Run log

#### Run summary

##### DATA OUTPUT

Estimated bases: **8.7 Gb**

Reads generated: **415.84 k**

Estimated N50: **47.16 kb**

##### BASECALLING

Reads called: **100%**

Reads called (min Q score: 9)

| Pass | Fail |
|---|---|
| 375.53 k | 40.22 k |

Bases called (min Q score: 9)

| Pass | Fail |
|---|---|
| 7.75 Gb | 640.51 Mb |

##### RUN DURATION

Run time

99 hrs 1 min / 99 hrs 1 min

Elapsed time ■ Run limit □

Run status

***FINISHED*** · *Target runtime has been reached*

View unit abbreviations used in this report

#### Run configuration

##### RUN SETUP

| | |
|---|---|
| Flow cell type | FLO-PRO114M |
| Flow cell type alias | FLO-PRO114M |
| Flow cell ID | PBE41021 |
| Kit type | SQK-ULK114 |

##### RUN SETTINGS

| | |
|---|---|
| Run limit | 96 hrs |
| Pore scan freq. | 1.5 hrs |
| Reserved pores | On |
| Basecalling | High-accuracy model 400bps |
| Modified basecalling | On |
| Modifications | • 5hmC & 5mC in all contexts |
| Min Q score | 9 |

##### DATA OUTPUT SETTINGS

| | |
|---|---|
| FAST5 output | Off |
| FASTQ data output | One file per hour |
| POD5 data output | One file per hour, or 500000000 bases per batch |
| BAM file output | On |
| BAM data output | One file per hour |
| Bulk file output | Off |
| Data location | /Library/MinKNOW/data/ElymusCiliaris-YM2/ECI_YM2/20250925_1154_P2S-02009-A_PBE41021_3d5de01d |

##### SOFTWARE VERSIONS

| | |
|---|---|
| MinKNOW | 25.05.12 |
| Bream | 8.5.4 |
| Configuration | 6.5.7 |
| Dorado | 7.9.8 |
| MinKNOW Core | 6.5.13 |

### Sequence output

#### READ LENGTHS · OUTLIERS REMOVED

The read length graph shows the total number of bases vs the read length. The longest 1% of strands are classified as outliers, and excluded to allow focus on the main body of data.

N50*: **47.16 kb**

% Basecalled: **100 %**

Basecalled | Estimated

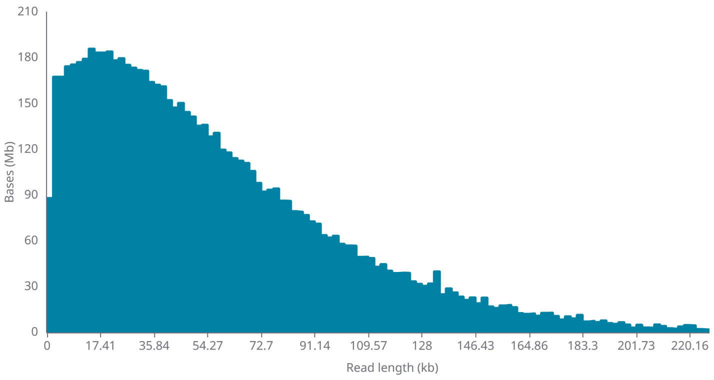


*N50 calculated from basecalled read length histogram.

#### OUTLIERS

The longest 1% of strands are classified as outliers, and aggregated into groups to show their relative amounts.

| Read length (kb) | Bases (Mb) |
|---|---|
| 237.568 - 368.64 | 26.92 |
| 368.64 - 499.712 | 0.37 |
| 499.712 - 630.784 | 0.56 |
| 630.784 - 761.856 | None |
| 761.856 - 786.432 | 0.78 |

#### CUMULATIVE OUTPUT

The cumulative output shows the total amount of bases or reads sequenced over time by your device.

##### Bases

###### Legend

- **Estimated**: Predicted total number of bases, prior to basecalling
- **Passed**: Bases equal to or above the quality score threshold.
- **Failed**: Bases below the quality score threshold.

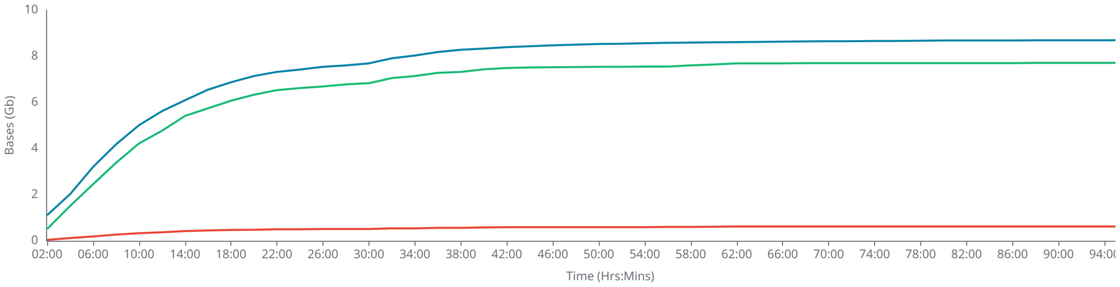


##### Reads

###### Legend

- **Total**: Total number of reads, including passed, failed and skipped.
- **Passed**: Reads equal to or above the quality score threshold.
- **Failed**: Reads below the quality score threshold.
- **Skipped**: Reads that will not be basecalled. Post run basecalling is possible.

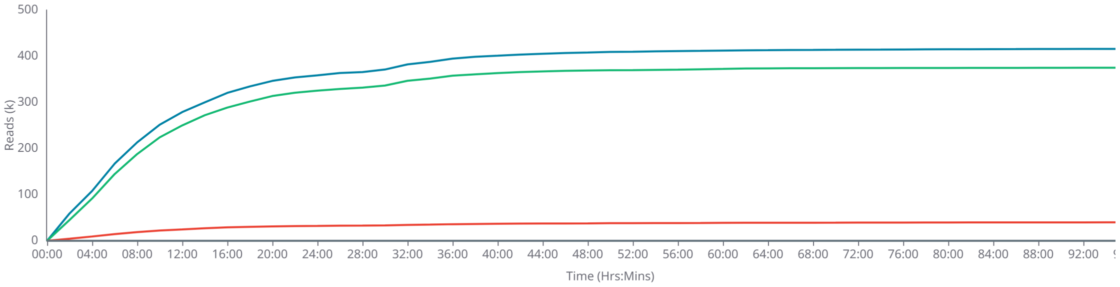

#### QUALITY SCORE

The quality score is calculated as basecalling is performed on your device. Reads that fall below the minimum value of 9 will be classified as failed reads. You can alter the accepted minimum quality score in MinKNOW.

##### Q score histogram

Passed simplex bases

Failed simplex bases

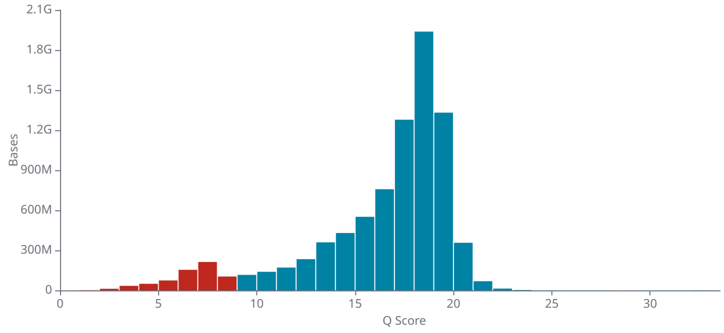


**Filters**

| | |
|---|---|
| Min Q score | 9 |

##### Q Score over time

**Legend**

**Mode** — The most frequent quality score of reads in the run.

**Spread** — The spread of quality scores, found by calculating full width half maximum.

**Min. quality score** — Minimum quality score to be accepted as a passed read.

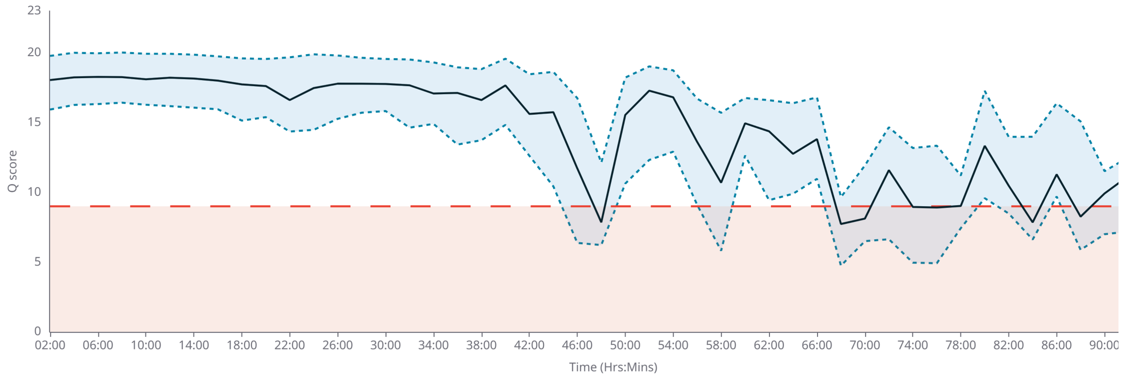


###### Troubleshooting

Quality score low

This can be due to the translocation speed being out of the accepted range, which can correlate to low quality scores. If you see that the translocation speed is out of the accepted range in the below graph, please see the Flow Cell refuelling page linked here for further troubleshooting.

### Run health

#### PORE ACTIVITY

The Pore activity graph shows the performance of your sample as it is being sequenced during a run.

☑ Show grouped

Legend

- **Sequencing** – Pore currently sequencing
- **Pore available** – Pore available for sequencing
- **Unavailable** – Pore currently unavailable for sequencing
- **Inactive** – Pore no longer suitable for further sequencing
- **Unclassified** – Pore status unknown

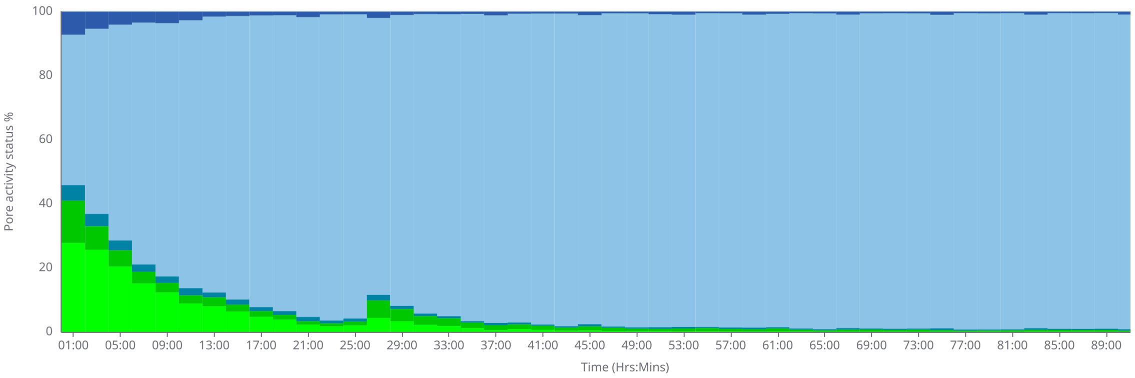


##### Troubleshooting

General

Some commonly seen issues are excess pores classified as Recovering, Open Pore, or Free Adapter. To find out what advice is applicable for your run, visit the user guide.

#### PORE SCAN

A Pore scan is performed at configurable time intervals to determine the current status of pores within channels on a Flow Cell. For this run a Pore scan is performed every 1.5 hrs.

Legend

- **Pore available** – Pore in channel available for sequencing
- **Reserved pore** – Pore in reserve, will return to available when required
- **Unavailable** – Pore inhibited from sequencing
- **Saturated** – Possible contamination in the sample
- **Zero** – No current is passing through this pore, possibly due to bubbles on the membrane
- **Inactive** – Pore no longer suitable for further sequencing

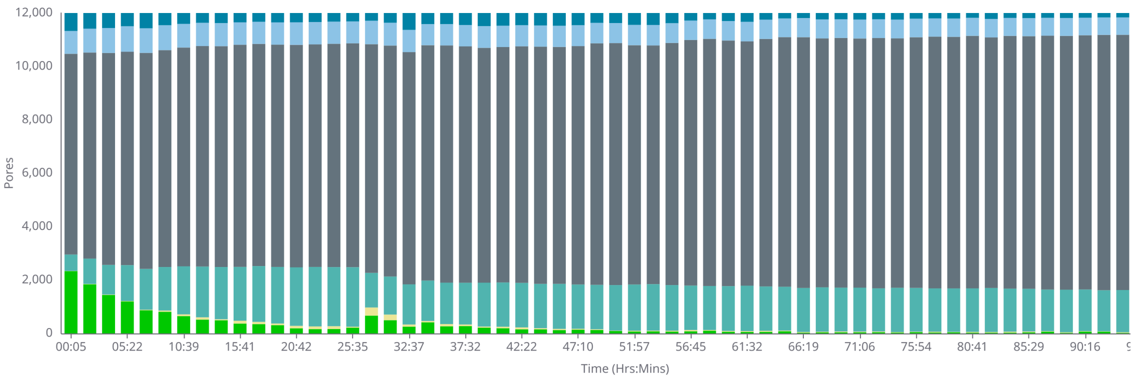


##### Troubleshooting

High proportion Unavailable

Possible contaminants in library blocking the pore. Consider using the Flow Cell Wash Kit, and reloading a library.

High proportion Inactive

If localised to one area of the Flow Cell, this could indicate that an air bubble has been introduced during the flushing/loading steps. If inactivity is spread across the Flow Cell this could be caused by improper loading of the library, please refer to the user guide for further support.

#### TRANSLOCATION SPEED

The translocation speed is the rate at which DNA/RNA travels through pores as it is being sequenced.

**Legend**

Median — 75% quartile — 25% quartile

Acceped range

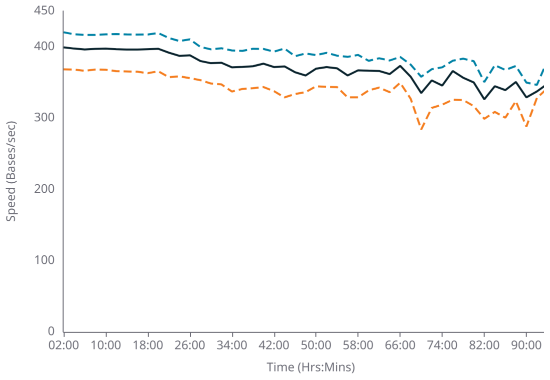


##### Troubleshooting

Low speed

Check that the Flow Cell is within the target temperature range.

Note

Low-quality and short reads are not included in this graph.

#### TEMPERATURE

The temperature of the Flow Cell over the run time.

**Legend**

Measured — Target

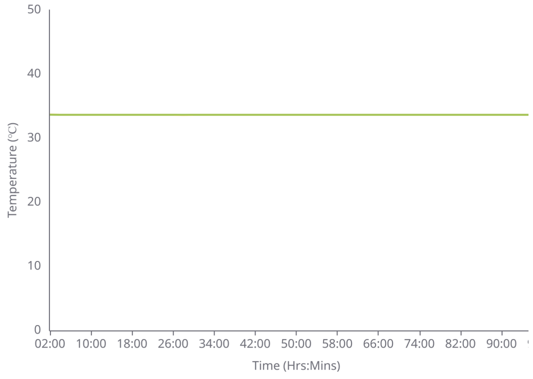


##### Troubleshooting

Out of range

Check that the Flow Cell is correctly seated and firmly pushed down into the device. Ensure ambient temperature is always within the specified range for your device in the user guide.
Air flow should be good but not excessive. Excessive amounts of cool air blowing on the device could prevent it from reaching target temperature.

#### Run log

##### SYSTEM MESSAGES

System messages are a record of the events that occurred in the time covered by this report.

###### Errors

###### Warnings

###### Events

#### Unit abbreviations

| | |
|---|---|
| Byte | B |
| Kilobyte | KB |
| Megabyte | MB |
| Gigabyte | GB |

| | |
|---|---|
| Base | b |
| Kilobase | kb |
| Megabase | Mb |
| Gigabase | Gb |
| Terabase | Tb |

| | |
|---|---|
| Minutes | mins |
| Hours | hrs |

Generated using Run Report Template v6.5.0
