## Supplementary material for "Chloroplast Genome Evolution, Heteroplasmy, and Inverted Repeat Dynamics in the *Elymus* Complex (Triticeae, Poaceae): Insights from Single-Molecule Sequencing of *Elymus ciliaris* and Comparative Analysis of St-Genome Lineages": Karimi_Elymus_SupplementaryFigureS1

**Supplementary Figure S1.** Gene map of the *Elymus ciliaris* chloroplast genome Variant 1R (135,008bp). Genes (total of 127) inside and outside the outer circle are transcribed clockwise and counterclockwise, respectively. Colors indicate functional groups: photosynthesis-related genes (green), transcription/translation-related genes (orange), rRNA genes (red), tRNA genes (blue). Dark gray shading inner circle represents GC content (average 38.3% GC); light gray represents AT content. IR, inverted repeat; LSC, large single copy; SSC, small single copy. See Fig. 2 for Variant 1F (135,004bp).

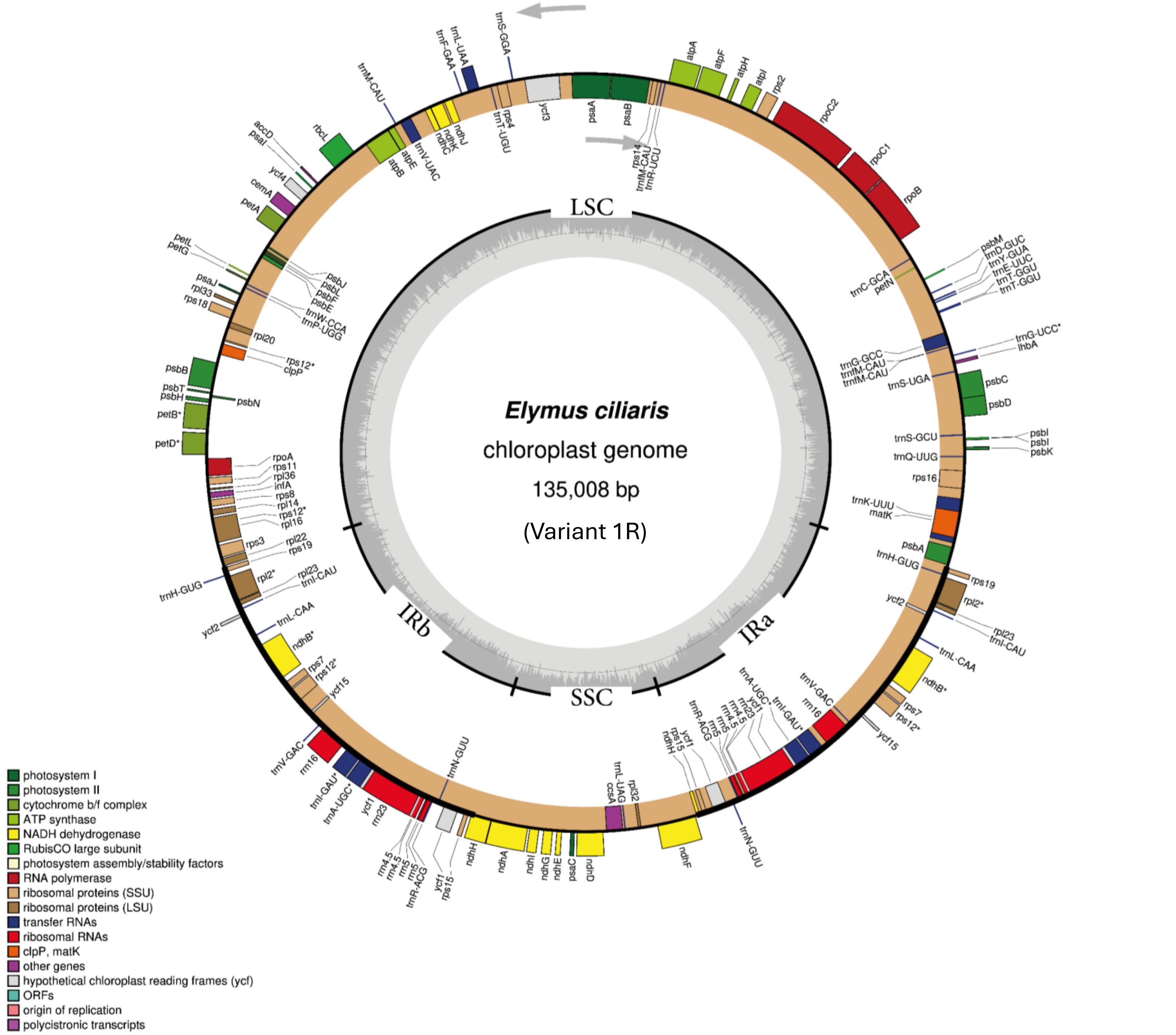
