## Supplementary material for "Chloroplast Genome Evolution, Heteroplasmy, and Inverted Repeat Dynamics in the *Elymus* Complex (Triticeae, Poaceae): Insights from Single-Molecule Sequencing of *Elymus ciliaris* and Comparative Analysis of St-Genome Lineages": Karimi_Elymus_SupplementaryFigureS2

**Figure S2**.Mauve alignments comparing chloroplasts of the *Elymus s.l.* of all species analysed showing nucleotide polymorphisms (red), but no notable interspecific rearrangements while reflecting small differences in length (vertical lines). Notably, no large-scale rearrangements were detected. Position in bp shown. A selection of genome alignments are shown in Figure 4.

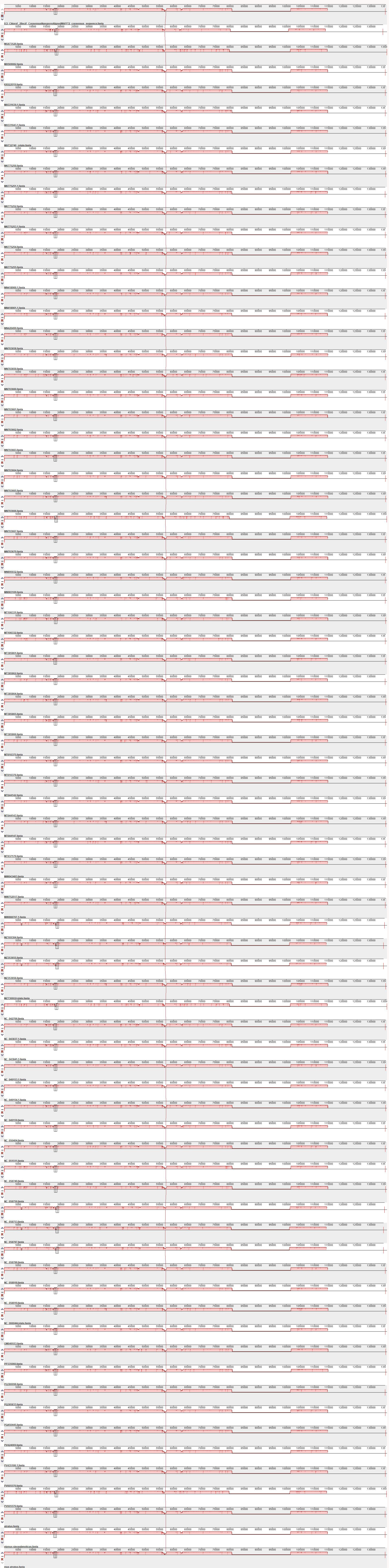
